# Loss of PRICKLE1 leads to abnormal endometrial epithelial architecture, decreased embryo implantation, and reduced fertility in mice

**DOI:** 10.1101/2024.08.06.605120

**Authors:** Emily R Roberts, Aishwarya V Bhurke, Sornakala Ganeshkumar, Sumedha Gunewardena, Ripla Arora, Vargheese M Chennthukuzhi

**Affiliations:** Department of Cell Biology and Physiology, University of Kansas Medical Center, Kansas City, KS, USA, 66160; Department of Obstetrics, Gynecology, and Reproductive Biology, Michigan State University, Grand Rapids, Michigan, USA, 49503; Institute for Quantitative Health Science and Engineering, Michigan State University, East Lansing, Michigan, USA, 48824; Department of Biostatistics, University of Kansas Medical Center, Kansas City, KS, USA, 66160

**Keywords:** Prickle1, Wnt/PCP, implantation, epithelial polarity, EMT

## Abstract

Successful embryo implantation requires coordinated changes in the uterine luminal epithelium, including structural adaptations, apical-basal polarity shifts, intrauterine fluid resorption, and cellular communication. Planar cell polarity (PCP) proteins, essential for cell organization, are understudied in the context of uterine physiology and implantation. PRICKLE proteins, components of PCP, are suggested to play critical roles in epithelial polarization and tissue morphogenesis. However, their function in the polarized unicellular layer of endometrial epithelium, which supports embryo implantation, is unknown. We developed an endometrial epithelial-specific knockout (cKO) of mouse *Prickle1* using *Lactoferrin-iCre* to investigate its’s role in uterine physiology. *Prickle1* ablation in the endometrial epithelium of mice resulted in decreased embryo implantation by gestational day 4.5 leading to lower fertility. Three-dimensional imaging of the uterus revealed abnormal luminal folding, impaired luminal closure, and altered glandular length in mutant uteri. Additionally, we observed decreased aquaporin-2 expression, disrupted cellular architecture, and altered E-Cadherin expression and localization in the mutant uterine epithelium. Evidence of epithelial-mesenchymal transition (EMT) was found within luminal epithelial cells, further linking PRICKLE1 loss to uterine pathologies. Furthermore, altered polarity of cell division leading to incomplete cytokinesis and increase in binuclear or multinucleated cells suggests a crucial role for PRICKLE1 in the maintenance of epithelial architecture. Our findings highlight PRICKLE1’s critical role in the PCP pathway within the uterus, revealing its importance in the molecular and cellular responses essential for successful pregnancy and fertility.

**Significance Statement:** Conservative cell division is essential to maintain apical-basal polarity and proper epithelial function in the uterus. Wnt/ Planar cell polarity signaling molecules are hypothesized to provide the spatial cues to organize unicellular, 2-dimensional sheet of epithelium in a plane orthogonal to the apical-basal polarity. Conditional ablation of *Prickle1*, a crucial Wnt/ PCP gene, in mouse uterine epithelium results in aberrant expression of epithelial cadherin, altered plane of cell division, incomplete cytokinesis leading to binucleated/ multinucleated cells, epithelial – mesenchymal transition, and defective implantation. Role of *Prickle1* in maintaining symmetric uterine epithelial cell division and tissue architecture is unique among Wnt/PCP genes, including previously described mouse models for *Vangl2, Ror2,* and *Wnt5a*.

**Classification:** Biological Sciences (Major) Cell Biology (Minor), Physiology (Minor)

*Graphical Abstract:* Conditional ablation of *Prickle1*, a crucial Wnt/ PCP gene, in mouse uterine epithelium results in altered plane of cell division and incomplete cytokinesis leading to binucleated/multinucleated cells, epithelial – mesenchymal transition, altered gland length, and defective implantation. Some images adapted from BioRender.com (2024).

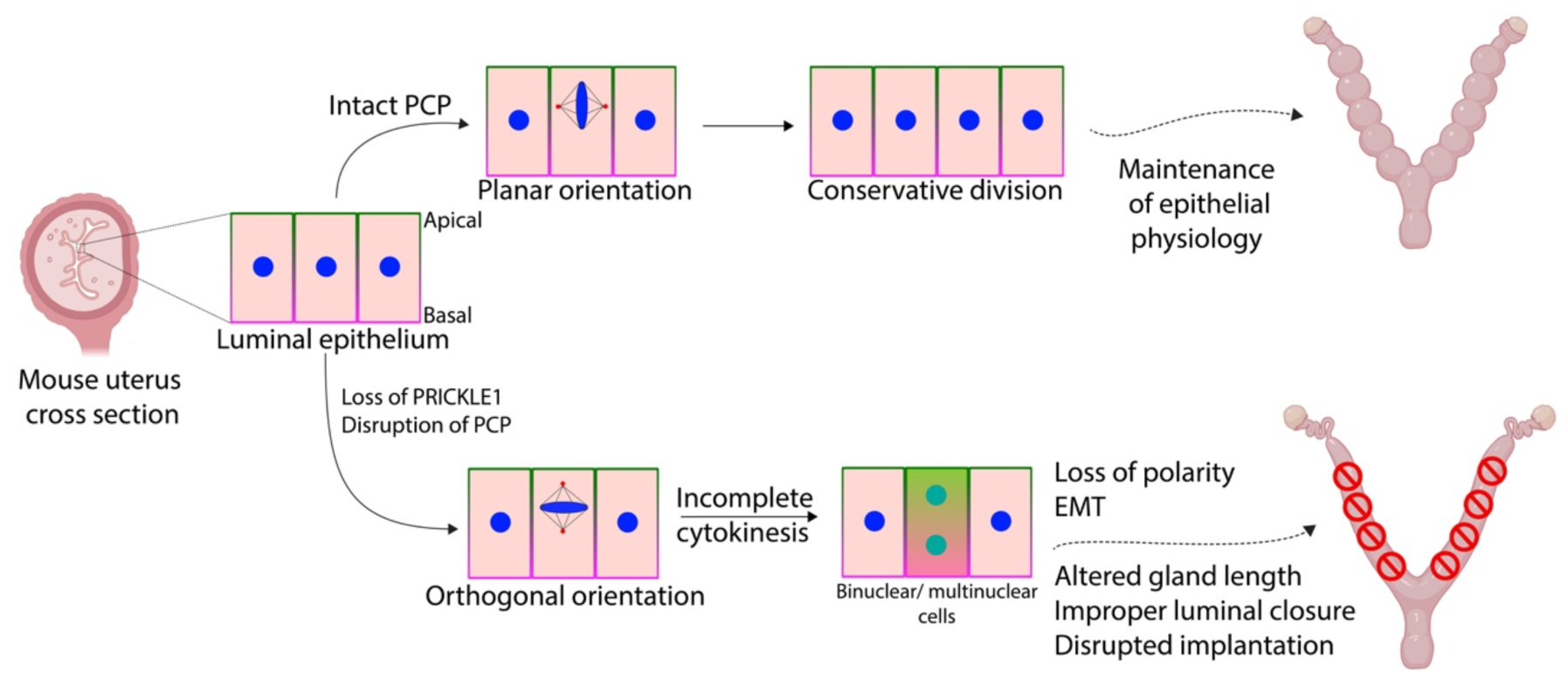

## Introduction

The uterine luminal epithelium is the primary maternal contact for the implanting embryo and several steps must coincide for successful embryo implantation to occur (1). These steps include precise timing of luminal epithelial structural changes (2, 3), appropriate alterations to apical-basal polarity (4, 5), intrauterine fluid resorption (6), and cellular communication between endometrial and uterine immune cells (7, 8). However, how the uterus transiently transforms into this receptive state for proper embryo implantation is poorly understood.

The luminal epithelium must undergo several morphological changes to prepare for an implanting embryo, and this term has been previously coined plasma membrane transformation (PMT) (5). During this process, the timing of this transition and the associated alteration to the cell structure is crucial to its success (4, 9–11). Major components of this process are specifically associated with planar cell polarity (PCP), a crucial component for proper epithelial layer formation and epithelial cell division in a specified plane of tissue (12). Some studies have investigated the chronology of these events utilizing the defective implantation of *Wnt5a* mutant mice (13). Moreover, recent work in uterine deletion of Van-Gogh-like 2 (*Vangl2*) has shown the importance of PCP and associated proteins in embryo implantation chamber formation and successful implantation in mice (14). However, the role of PCP signaling in the uterine epithelial morphogenesis, and its function during implantation is still widely not understood.

Planar cell polarity proteins, first identified in drosophila (15–18), provide vectorial information for cell organization and migration across a wide field. However, of the six main PCP proteins, PRICKLE has been largely understudied and particularly unstudied in PCP within the context of uterine physiology and the process of embryo implantation. The mouse genome contains four *Prickle* homologs, where *Prickle1* and *Prickle2* are the most closely related to drosophila *Prickle* gene. While the role of PRICKLE1 and PRICKLE2 has been previously studied in embryonic AB polarity (19, 20), the early embryonic lethality of their conventional knockouts has limited our knowledge of their role in late-stage tissue development (21, 22). PRICKLE1 exerts multifaceted roles in cellular polarization and tissue morphogenesis and has been implicated in regulating the non-canonical Wnt/PCP pathway (23). PRICKLE1 has yet to be studied in the context of the endometrial epithelium and its role in PCP maintenance within the uterus.

To explore the PCP-related function of PRICKLE1 in uterine physiology and embryo implantation, we developed an endometrial epithelial conditional knockout (cKO) of *Prickle1* using a *Lactoferrin-icre*. Our results demonstrate that the loss of PRICKLE1 in the endometrial epithelium leads to decreased fertility and embryo implantation defect as early as gestational day 4.5. In addition, three-dimensional imaging demonstrates aberrant luminal folding, impaired luminal closure, and altered glandular structure in mutant uteri at gestational day 4.5. Further analysis demonstrated decreased aquaporin-2 (AQP2) expression in mutant mouse uteri during diestrus. Moreover, we provide evidence for altered PCP via dysregulation of cellular architecture, altered E-Cadherin expression and localization, and aberrant gene expression in non-pregnant and gestational day 3.5 mutant mice. Additionally, the loss of PRICKLE1 in endometrial epithelium shows evidence of epithelial-to-mesenchymal transition within luminal epithelial cells of non-pregnant mice, providing a connection to uterine pathologies. Finally, the loss of PRICKLE1 in endometrial epithelium shows alterations to the plane of cellular division, resulting in asymmetric cell division, increased multinucleated cells and providing a connection to the evidence of EMT. PRICKLE1 involvement in the PCP pathway within the uterus and during embryo implantation unveils a complex interplay between molecular cues and cellular responses vital for successful pregnancy establishment.

## Results

### cKO of *Prickle1* in Mouse Endometrial Epithelium Leads to Alterations in Luminal Folding, Intrauterine Fluid Resorption, and Decreased Embryo Implantation

Using *Prickle1^f/+^*ES cells, we developed *Prickle1^f/f^*mice and utilized *lactoferrin-icre* (24) to conditionally ablate *Prickle1* in the mouse endometrial epithelium (Figure 1 A-D).

**Figure 1.**
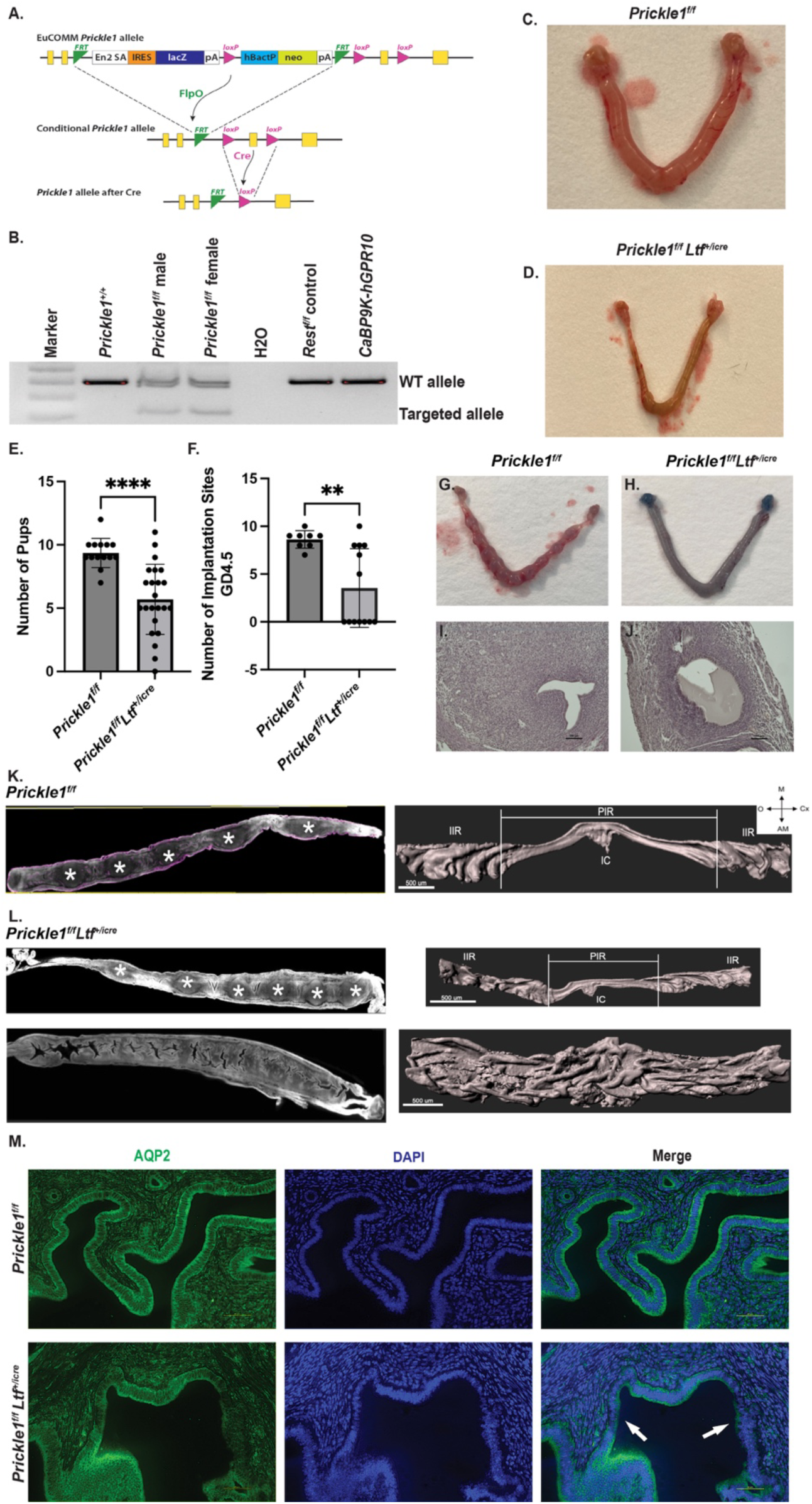
Loss of *Prickle1* under the *Ltf* promoter displays alterations to luminal folding and decreased embryo implantation. (A) *Prickle1^f/f^* targeting construct. (B) PCR genotyping of *Prickle1^f/f^*mice (lanes 3 and 4), *Prickle1* heterozygous mice (lane 1), and control mice (lanes 4, 5, and 6). (C,D) Representative images showing altered morphology of uterus in 6-month-old *Prickle1^f/f^ Ltf^+/icre^* cKO mouse (D) compared to control (C) mouse during diestrus. (E) Fertility assessment in control (n=14) and *Prickle1^f/f^ Ltf^+/icre^* cKO (n=23) 6-month-old mice showing decreased fertility in *Prickle1^f/f^ Ltf^+/icre^* cKO. (F) Implantation assessment at gestational day 4.5 of control (n=8) and *Prickle1^f/f^ Ltf^+/icre^* cKO (n=13) 6-month-old mice indicated two distinct groups of mutants. **P<0.05, ****P<0.0001. (G-J) Representative images showing implantation sites with corresponding H&E stain in 6-month-old *Prickle1^f/f^ Ltf^+/icre^*cKO mouse (H and J) compared to control (G and I) mouse at gestational day 4.5 showing abnormal fluid accumulation at proposed embryo site of *Prickle1^f/f^ Ltf^+/icre^* cKO. (Scale bar, 100 μm) (K,L) Optical z slice showing one horn of *Prickle1^f/f^ Ltf^+/icre^* cKO (L) and control uteri (K) with gestational day 4.5 embryos (*) with corresponding 3-dimensional surface models showing aberrant luminal folding in the mutant with no embryos present. (Scale bar, 500 μm) (PIR, peri-implantation region; IIR, inter-implantation region; IC, implantation chamber; M, mesometrial pole; AM, anti-mesometrial pole; O, ovary; Cx, cervix). (M) Uterine cross-sections of 6-month-old *Prickle1^f/f^ Ltf^+/icre^* cKO and control mice in diestrus stained with AQP2 (green) and DAPI (blue). Arrows indicate areas of low AQP2 expression. (Scale bar, 50 μm).

To investigate the impact that the loss of PRICKLE1 within the endometrial epithelium had on fertility, *Prickle1^f/f^ Ltf^+/icre^* cKO female mice were bred with C57BL6 WT male mice and the number of live birth pups was tracked following a positive vaginal plug formation. These fertility trials indicated that *Prickle1^f/f^ Ltf^+/icre^* cKO mice have a reduced litter size of 5.70 pups per litter compared to 9.36 pups per litter in *Prickle1^f/f^*control mice (p<0.0001) (Figure 1 E). However, a wide range of litter sizes was observed within the cKO groups, indicating that *Prickle1^f/f^ Ltf^+/icre^* cKO mice may have a range in the severity of knockout. Therefore, implantation studies were performed at gestational day (GD) 4.5 following breeding with C57BL6 WT male mice to investigate the timing of proposed improper embryo implantation. These studies demonstrated that *Prickle1^f/f^ Ltf^+/icre^* cKO mice show two distinct mutant groups: one group with implantation site number similar to that of control mice and one group with no implantation sites present at GD4.5 (Figure 1 F-J), potentially resulting from the incomplete penetrance of *iCre* expression (25). The results showing a range of live births from *Prickle1^f/f^ Ltf^+/icre^*cKO females (Figure 1 E) indicate further loss of embryos during pregnancy, in addition to the implantation defects.

Past studies have examined the importance of luminal folding during peri-implantation to the success of embryo implantation and decidualization (4, 26). To further investigate how the mutant’s uterine structures and luminal folding may differ from control mice, 3-dimensional imaging of full uterine horns with GD4.5 implantation sites was performed (Figure 1 K and L). This imaging confirmed the presence of two types of mutants where one set of *Prickle1^f/f^ Ltf^+/icre^* uteri have normal embryos present, normal luminal closure, and normal transverse luminal folding and decidual sites comparable to control mice, while the other set of *Prickle1^f/f^ Ltf^+/icre^* uteri have no embryos present, a completely open lumen, aberrant uterine folding, and no decidua. These more severe mutant mice demonstrate a super-folded luminal nature, indicating improper luminal folding and closure during peri-implantation.

Several studies have demonstrated the involvement of aquaporins in the luminal closure and their dynamic regulation during implantation (27–29). Immunostaining of non-pregnant mice at diestrus as well as GD3.5 *Prickle1^f/f^Ltf^+/icre^*uteri demonstrated areas of altered aquaporin 2 (AQP2) expression as compared to control uteri (Figure 1 M, SI Appendix, Figure S1). Moreover, a receptive uterus is marked by the downregulation of *Muc-1* in the luminal epithelium and the cessation of epithelial cell proliferation prior to implantation (30–32). It has been well established that the persistent expression of *Muc-1* and stromal growth factors impede implantation (33). Here, we see an increase of fibroblast growth factors *Fgf1, Fgf2, Fgf7, Fgf9, Fgf10,* and *Fgf18* in addition to *Muc-1* mRNA expression in the *Prickle1^f/f^ Ltf^+/icre^* cKO when compared to control (SI Appendix, Figure S2). In addition, we see altered expression of *Pgr* and *Esr1* across all estrus cycle stages (SI Appendix, Figure S3) and increased *Pgr, Esr1, and Muc-1* at GD4.5 (SI Appendix, Figure S4), indicating aberrant steroid hormone signaling and altered epithelial-stromal crosstalk. Hormone assessment of cKO in diestrus was comparable to controls (SI Appendix, Figure S5).

While evidence of luminal epithelial changes was seen within the *Prickle1^f/f^ Ltf^+/icre^* uteri, the glandular epithelium remained largely intact but showed altered gland length in 3D (Figure 2 A-D). At GD4.5, as observed previously, glands were tubular and branched in control uteri with a median gland length of 353.62 μm (13, 34). While the glands of the *Prickle1*^f/f^ *Ltf*^+/icre^ cKO mice uteri appeared similar to control uteri with respect to their tubular and branched nature, the glands of mutant uteri that displayed open lumens were significantly shorter in length (median = 292.87 μm) and glands of the mutant uteri that displayed closed lumens showed an increase in length (median = 440.92 μm) (Figure 2 C and D).

**Figure 2.**
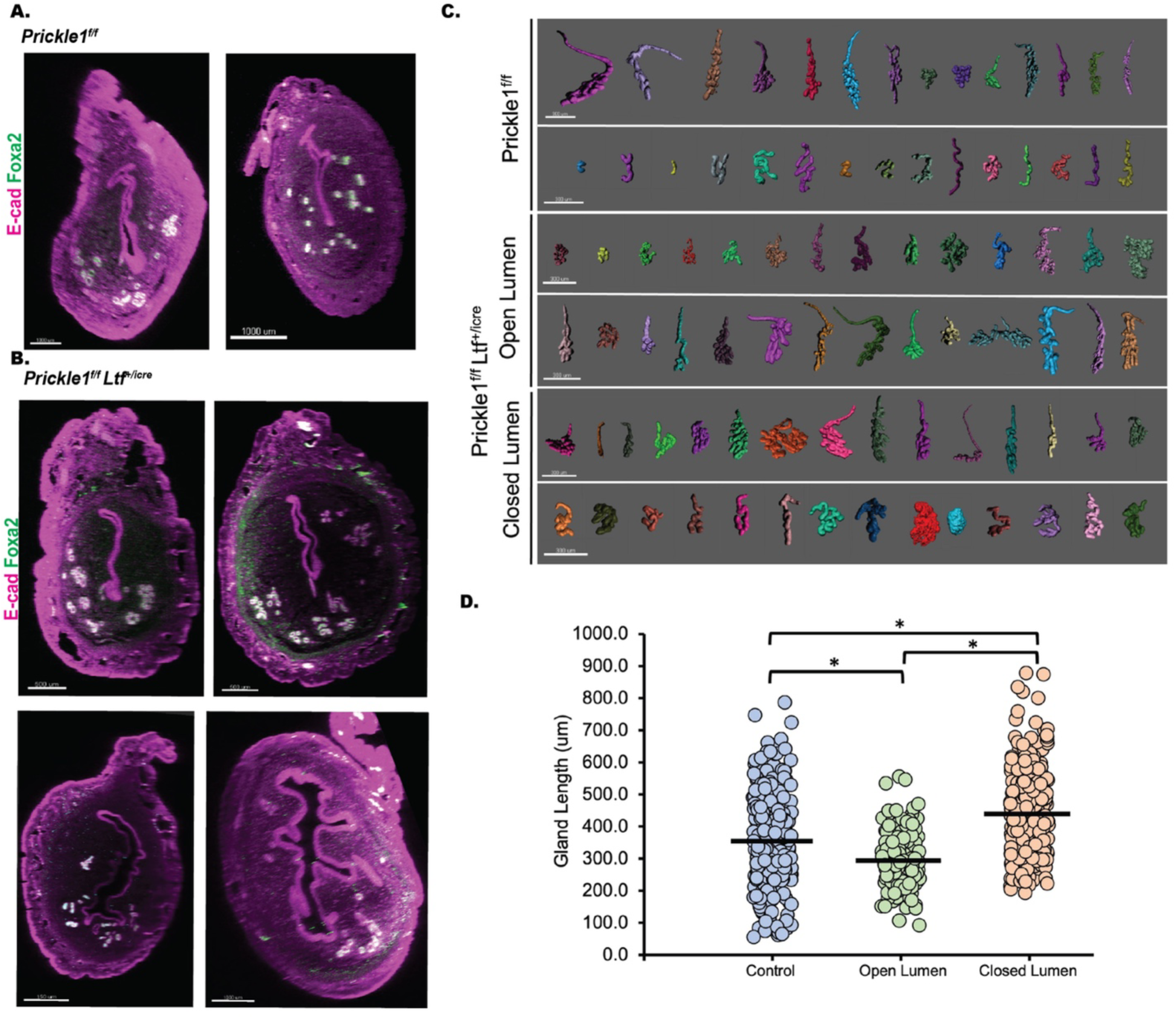
Loss of *Prickle1* under the *Ltf* promoter does not alter glandular structure at gestational day 4.5. (A,B) Transverse slices of 6-month-old *Prickle1^f/f^ Ltf^+/icre^* cKO (B) compared to control (A) mice at gestational day 4.5 stained with epithelial marker E-cadherin (purple) and glandular marker FOXA2 (green). (Scale bars, 500 μm (controls), 1000 μm (*Prickle1^f/f^ Ltf^+/icre^* section 1 and 2), 150 μm (*Prickle1^f/f^ Ltf^+/icre^* section 3), and 1000 μm (*Prickle1^f/f^ Ltf^+/icre^* section 4)). (C) Representative 3D reconstructions of glands of 6-month-old *Prickle1^f/f^ Ltf^+/icre^* cKO compared to control mice at gestational day 4.5. (Scale bars, 300 μm). (D) Quantitative analysis of average gland length measurements (each dot represents one gland). 250-470 glands analyzed per mouse. Data analyzed using Kruskal-Wallis test with Dunn’s multiple comparisons. *P<0.05.

### cKO of *Prickle1* in Mouse Endometrial Epithelium Leads to Changes in Epithelial Architecture and Promotes EMT

To further elucidate morphological alterations, cross-sections of *Prickle1^f/f^ Ltf^+/icre^* uteri were examined and revealed prominent areas with aberrant epithelial cell shape and presence of multinucleated cells (Figure 3 A, B, and E, and SI Appendix, Figure S6). Moreover, immunostaining for PCP proteins showed increased overall expression of E-cadherin with increased basal expression as compared to lateral and apical expression in the control uteri across all cycle stages of estrus (Figure 3 C and D and SI Appendix, Figure S7), providing evidence for PCP disruption (35). Furthermore, *Prickle1^f/f^ Ltf^+/icre^* uteri showed increased overall cytokeratin 7 (KRT7) expression (SI Appendix, Figure S8) and increased SOX9 nuclear expression (56.1%) compared to control uteri (31.4%) (Figure 3 F-H), a marker of endometrial hyperplasia and endometriosis (36, 37), demonstrating that the loss of PRICKLE1 in the endometrial epithelium leads to PCP disruption. In addition, these results were consistent at GD3.5 (SI Appendix, Figure S1), indicating that this altered PCP expression is present both during the cycle and just prior to implantation.

**Figure 3.**
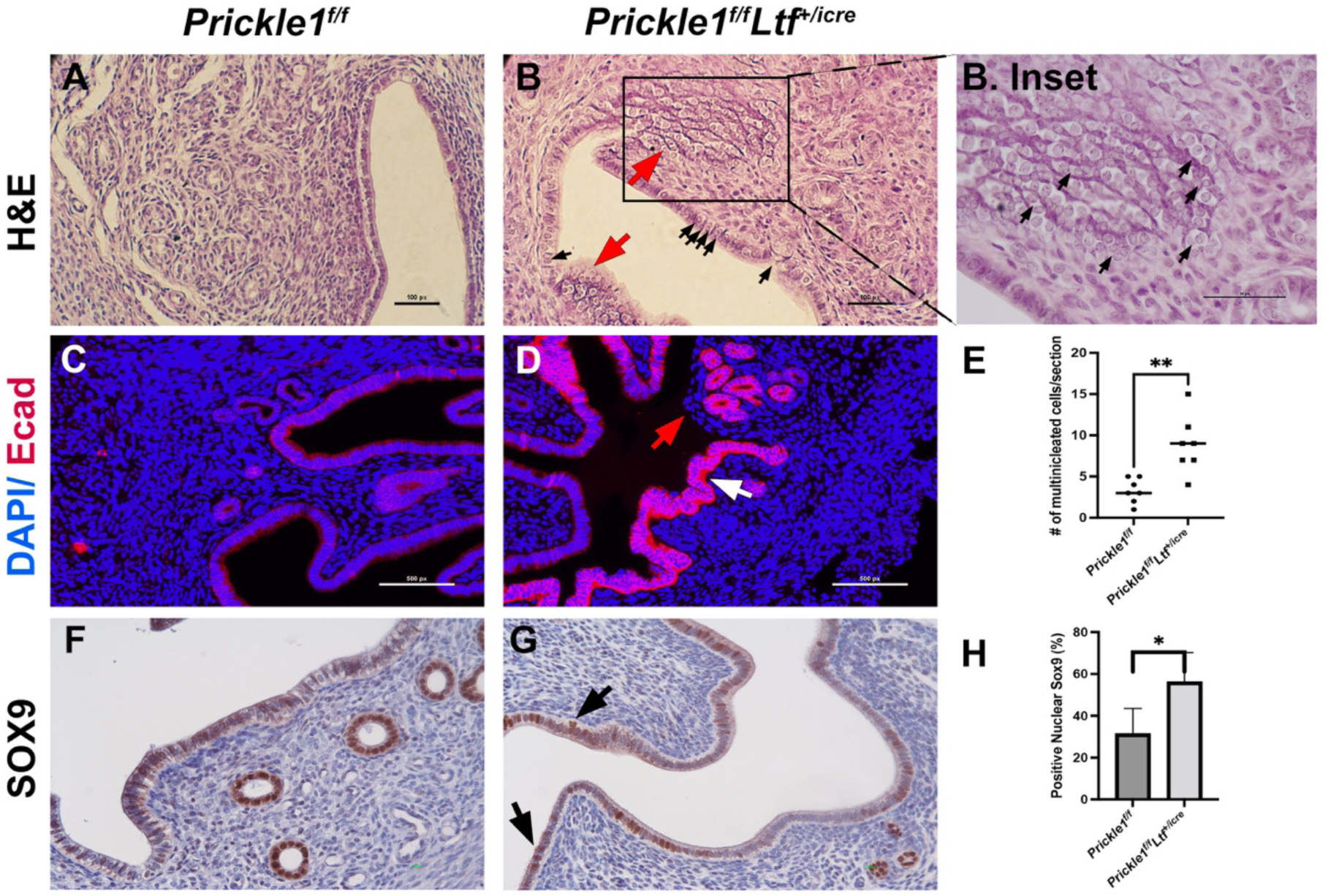
Loss of *Prickle1* under the *Ltf* promoter alters cellular structure and dysregulates expression of Wnt/PCP proteins. (A,B) H&E stain of 6-month-old *Prickle1^f/f^ Ltf^+/icre^*cKO (B) compared to control (A) in diestrus. Red arrows indicate areas of hyperproliferation and rounded epithelial shape. Black arrows indicate multinucleated cells. (Scale bars, 100 μm) (B inset) Callout of *Prickle1^f/f^ Ltf^+/icre^* cKO highlights area of multinucleated and spindle shaped cells. Arrows indicate multinucleated and spindle shaped cells. (Scale bars, 50 μm). (C,D) Immunofluorescence stain of 6-month-old *Prickle1^f/f^ Ltf^+/icre^* cKO (C) compared to control (D) in diestrus for epithelial marker E-cadherin (red) and DAPI (blue) indicating high baso-lateral and overall E-cadherin expression in mutant. Red arrow indicates area of missing epithelium and black arrow indicates high basal expression of E-cadherin. (Scale bars, 500 μm). (E) Quantification of multinucleated cells in 6-month-old *Prickle1^f/f^ Ltf^+/icre^*cKO (n=7) compared to control (n=7) in diestrus. ** P<0.005. (F,G) Immunohistochemistry stain of 6-month-old *Prickle1^f/f^ Ltf^+/icre^* cKO (G) compared to control (F) in diestrus for SOX9. Arrows indicate areas of nuclear expression and multinucleated cells in the mutant. (Scale bars, 10 μm). (H) Quantification of positive SOX9 nuclear staining in 6-month-old *Prickle1^f/f^ Ltf^+/icre^* cKO (n=5) compared to control (n=5) in diestrus. *P<0.05.

While alterations to structure and function were seen from our analysis of the uterus as a whole tissue, further understanding of individual cellular changes was pertinent. Therefore, to study the impact the loss of PRICKLE1 in the endometrial epithelium has on the whole uterus by cell type, single-cell RNA (scRNA) sequencing was performed on 6-month-old *Prickle1^f/f^ Ltf^+/icre^* and *Prickle1^f/f^* mice in diestrus (GSE272552). Results from Seurat single-cell RNA-sequencing data analysis (38), which passed the quality-control markers before being analyzed (SI Appendix, Figure S9), identified 12 clusters in the uterus based on their expression profiles (Figure 4 A). Cell types were determined via gene expression profiles using SingleR software (39) and expert curation. Based on conserved gene expression, clusters were identified as stromal (clusters 0, 4, and 5) and epithelial (clusters 3 and 8) (SI Appendix, Figures S10 and S11).(40)

**Figure 4.**
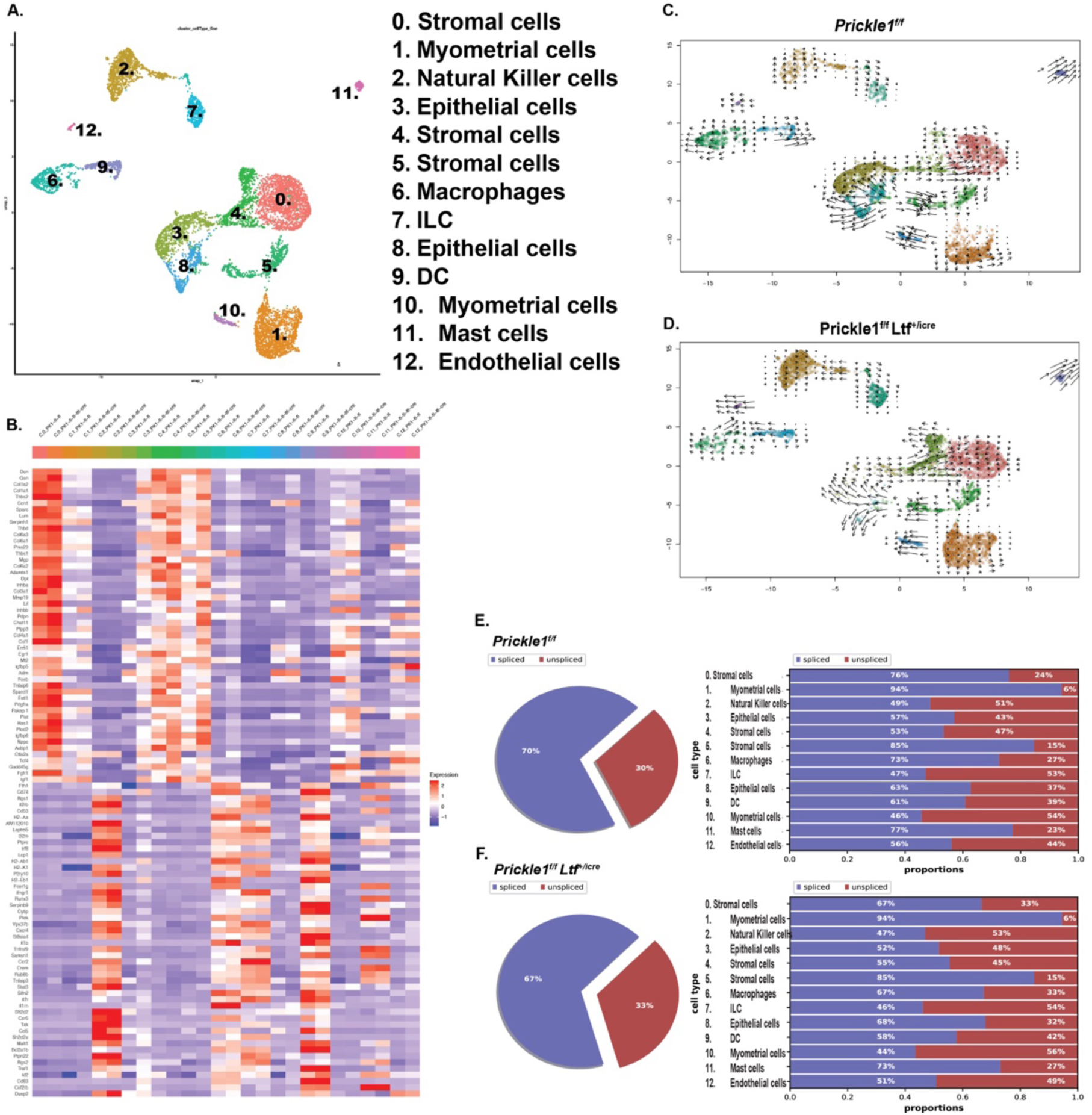
Loss of *Prickle1* under the *Ltf* promoter promotes increased expression of collagens, Wnt/PCP, and EMT markers in stromal clusters (0, 4, and 5). (A) SingleR program predictions for cell clusters present in control and *Prickle1^f/f^ Ltf^+/icre^*mice uteri shown by a UMAP plot. Cell clusters were given a number between 0-12. (B) Top TCA features (average cluster expression) of genes shown via heatmap showing upregulation of ECM components, Wnt/PCP, and EMT markers in stromal fibroblasts (clusters 0, 4, and 5) in *Prickle1^f/f^ Ltf^+/icre^* mice uteri. (C,D) Velocity RNA analysis embedding stream for control (C) and *Prickle1^f/f^ Ltf^+/icre^* (D) showing higher magnitude velocity in epithelial cells (cluster 3) of *Prickle1^f/f^ Ltf^+/icre^* mice. (E,F) Pie chart for entire sample and individual charts showing the percentage of spliced and unspliced RNA transcripts for each cluster for control (E) and *Prickle1^f/f^ Ltf^+/icre^* mice (F).

A χ^2^ test of homogeneity comparing *Prickle1^f/f^ Ltf^+/icre^* and *Prickle1^f/f^*scRNA sequencing counts revealed significant differences in populations of myometrial cells, epithelial cells, stromal cells, and natural killer cells (SI Appendix, Figure S12). Of particular interest was an over 11 times decrease in the number of epithelial cells in the *Prickle1^f/f^ Ltf^+/icre^* group as compared to the control, indicating a loss of epithelial-identified cells within the cKO group. A corresponding increase in stromal cells (clusters 0, 4, and 5) was observed. Moreover, there was 4 times increase in the number of natural killer cells present in the cKO when compared to the control.

Top PCA features for each cluster were identified and revealed increased REST target genes for clusters 0, 4, and 5 (stromal cells) (41) in addition to increased expression of various collagens (*Col1a1, Col1a2, Col6a1, Col6a3, Col6a2, Col4a1, Col3a1*) and known markers of epithelial-mesenchymal transition (EMT) (*Serpinh1, Ccn1, Lum, Mmp19, Lif, Igfbp6*) (42–47) within the *Prickle1^f/f^ Ltf^+/icre^* group compared to control (Figure 4 B and SI Appendix, Figure S13). In addition to PCP, PRICKLE1 is known to regulate the nuclear localization of REST (48).

In addition, decreased expression of *Cdh1, Muc1,* and *Tjp3* in epithelial cells while increased stromal expression of *Dvl1, Wnt5a, Tjp1, Tjp2,* and *Esr1* was seen (SI Appendix, Figure S14). These results indicate that prominent changes are occurring within the stromal cell population of the *Prickle1^f/f^ Ltf^+/icre^* cKO mice due to the loss of PRICKLE1 within the endometrial epithelium. Furthermore, the gene-expression profile of the epithelial cluster of the cKO showed similarities to cancer, organismal injury and abnormalities, reproductive system disease, and cell-to-cell signaling (Table 1-2, SI Appendix, Tables S1-S4) while stromal clusters of the cKO showed many of the same pathways in addition to inflammatory response, cellular movement, tissue morphology, and reproductive system development and function (Table 3, SI Appendix, Tables S5-S12). Moreover, the gene-expression profile of the natural killer cell cluster of the cKO showed connections to cancer, inflammatory disease, cell cycle, cellular growth and proliferation, and tissue development (SI Appendix, Tables S13-S15).

**Table 1.** Pathway analysis of predicted molecular and cellular functions associated with dysregulated genes in the epithelial cell population (cluster 3) of *Prickle1^f/f^ Ltf^+/icre^* cKO mice.

| **Molecular and Cellular Functions** | **P value** |
| --- | --- |
| Cellular movement | 4.82E-61 |
| Cell death and survival | 1.19E-47 |
| Cellular development | 4.12E-37 |
| Cellular growth and proliferation | 4.12E-37 |
| Cell-to-cell signaling and interaction | 1.22E-34 |

Function predictions made by IPA software based off genes which were found to be dysregulated in epithelial cells of 6-month-old *Prickle1<sup>ff</sup> Ltf<sup>+/fcre</sup>* mice cKO (n=3) in diestrus.

**Table 2.**
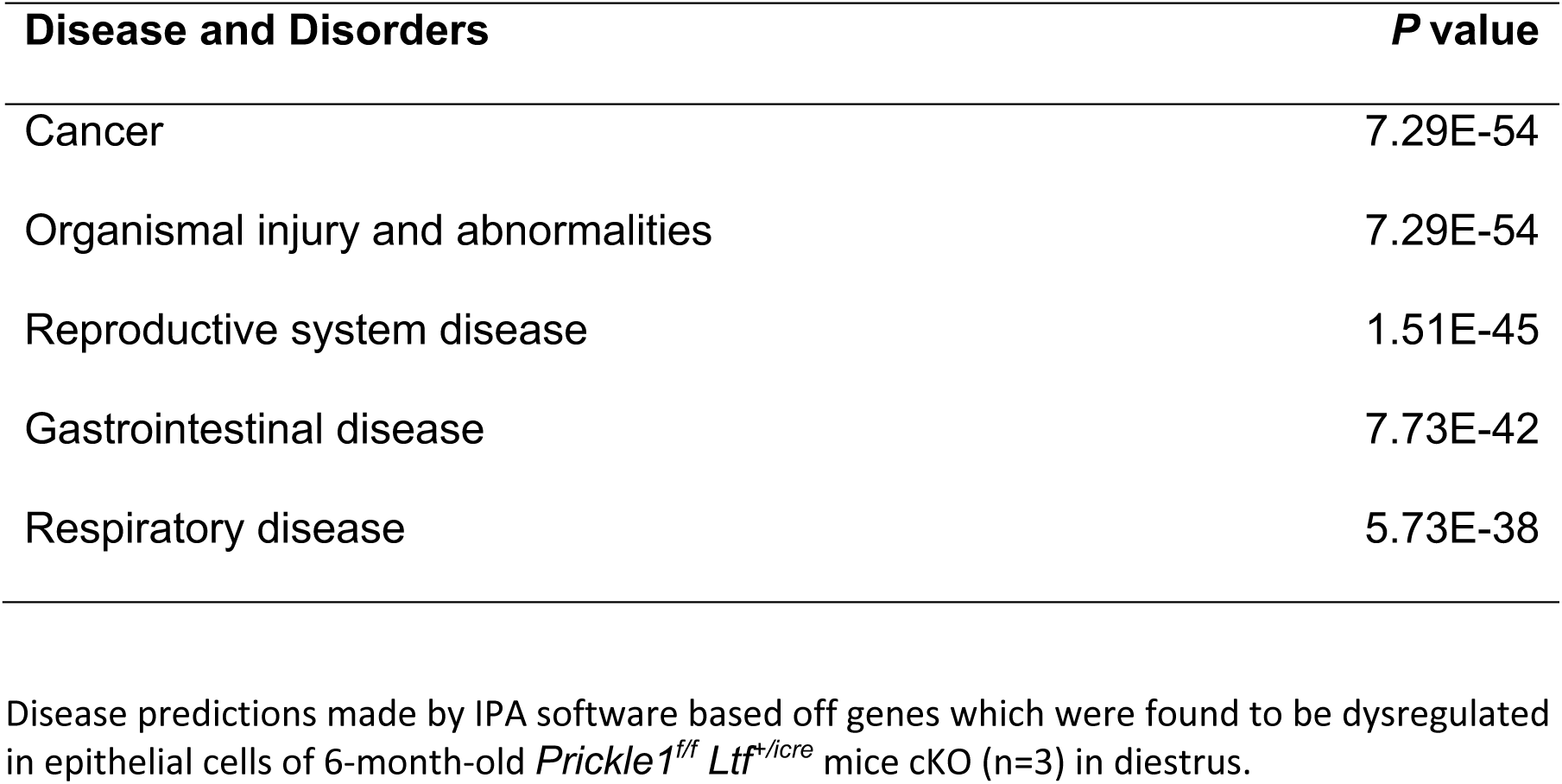
Pathway analysis of predicted disease and disorders associated with dysregulated genes in the epithelial cell population (cluster 3) of *Prickle1^f/f^ Ltf^+/icre^*cKO mice.

| Disease and Disorders | P value |
| --- | --- |
| Cancer | 7.29E-54 |
| Organismal injury and abnormalities | 7.29E-54 |
| Reproductive system disease | 1.51E-45 |
| Gastrointestinal disease | 7.73E-42 |
| Respiratory disease | 5.73E-38 |

Disease predictions made by IPA software based off genes which were found to be dysregulated in epithelial cells of 6-month-old *Prickle1<sup>ff</sup> Ltf<sup>+/iCre</sup>* mice cKO (n=3) in diestrus.

**Table 3.**
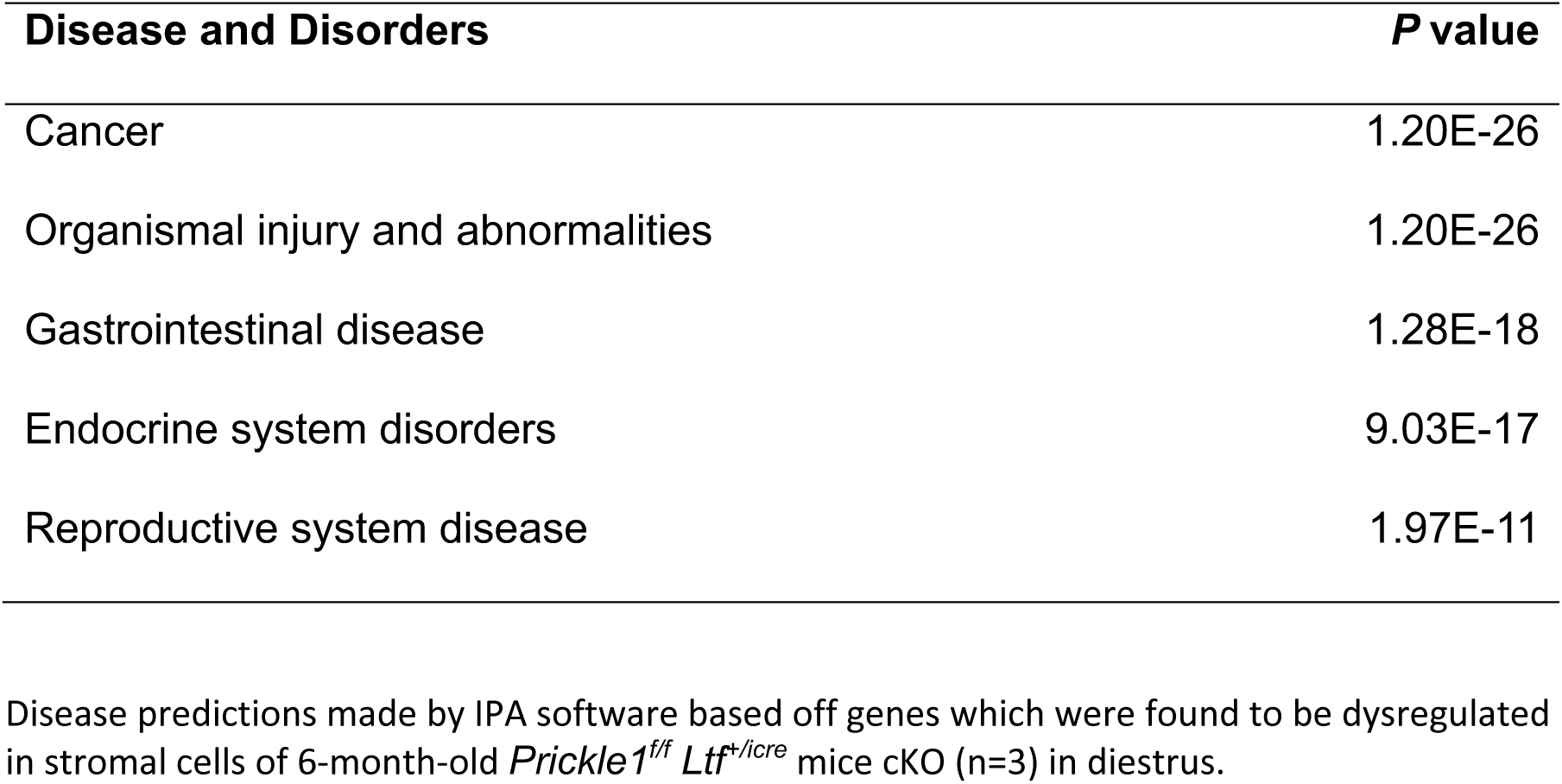
Pathway analysis of predicted disease and disorders associated with dysregulated genes in the stromal cell population (cluster 4) of *Prickle1^f/f^ Ltf^+/icre^*cKO mice.

| Disease and Disorders | P value |
| --- | --- |
| Cancer | 1.20E-26 |
| Organismal injury and abnormalities | 1.20E-26 |
| Gastrointestinal disease | 1.28E-18 |
| Endocrine system disorders | 9.03E-17 |
| Reproductive system disease | 1.97E-11 |

Disease predictions made by IPA software based off genes which were found to be dysregulated in stromal cells of 6-month-old *Prickle1<sup>ff</sup> Ltf<sup>+/icre</sup>* mice cKO (n=3) in diestrus.

In addition, a loss of cells within clusters 3 and 8 (epithelial cells) with a corresponding increase in cells within clusters 0, 4, and 5 (stromal cells) was observed within the *Prickle1^f/f^ Ltf^+/icre^*group as compared to the control group (Figure 4 C and D and SI Appendix, Figure S4). Given the previous evidence of altered gene expression and EMT markers in the stroma, we hypothesized that these epithelial groups were potentially undergoing EMT and re-clustering as stroma in the *Prickle1^f/f^ Ltf^+/icre^* cKO uteri. RNA velocity analysis was performed to identify high-dimensional vectors to predict the future state of the cells on a pseudo timescale (49) (Figure 4 E and F). Immediately, high velocity (as indicated by the higher ratio of unspliced: spliced RNA transcripts) was seen in clusters 0, 4, and 5 of the stroma of *Prickle1^f/f^ Ltf^+/icre^* cKO uteri as predicted, indicating large changes to the RNA transcripts of this group of cells in comparison to their control counterparts (Figure 4 C-F).

To further demonstrate that this reclassification of cells is due to EMT, RNA velocity analysis of 120 EMT hallmark genes revealed high velocity in clusters 3 and 8 (epithelial) to clusters 0, 4, and 5 (stroma) within the *Prickle1^f/f^ Ltf^+/icre^*cKO group when compared to the control group (SI Appendix 2, Figure A1A-L). These results provide evidence that *Prickle1* knockout in the endometrial epithelium promotes EMT within the epithelial cells and potentially reclassifies these cells as stromal cells during scRNA sequencing analysis.

To further validate the results of the RNA velocity analysis, immunostaining of EMT markers known to be upregulated in endometrial cancers (45, 50, 51) was performed on *Prickle1^f/f^ Ltf^+/icre^* uteri cross-sections. Increased expression of ZEB1 (SI Appendix, Figure S15 A), TWIST1 (SI Appendix, Figure S15 B), SNAI2/SLUG (SI Appendix, Figure S15 C), and Vimentin (SI Appendix, Figure S15 D) in both the endometrial epithelium and stroma of the cKO when compared to control uteri was observed, confirming evidence of increased EMT activity within the *Prickle1^f/f^ Ltf^+/icre^* cKO as previously seen by the scRNA sequencing analysis and providing further evidence for the loss of PRICKLE1 in promoting EMT with the endometrial epithelium.

### cKO of *Prickle1* in Mouse Endometrial Epithelium Leads to Altered Plane of Epithelial Cell Division

Further evaluation of the binuclear and multinucleated epithelial layer revealed alterations to the plane of cell division within the *Prickle1^f/f^ Ltf^+/icre^* uteri cross-sections with asymmetric epithelial cell division seen in the absence of PRICKLE1 (Figure 5 A-D). *Prickle1^f/f^ Ltf^+/icre^*uteri contained much higher rates of asymmetric epithelial cell division in comparison to controls (SI Appendix, Table S16). Moreover, higher levels of E-cadherin expression were present at basal and apical sides of the epithelium within the cKO uteri compared to controls, providing a potential mechanism for altered polarity and incomplete cell division (Figure 5 D and E). Finally, during scRNA data analysis, *Prickle1^f/f^ Ltf^+/icre^* cKO samples demonstrated a statistically higher occurrence of doublets (3.4%) as compared to control samples (2.5%) (P<0.05) (SI Appendix, Table S17), indicating a higher percentage of multinucleated cells present within the cKO samples, while assessment of apoptosis did not show any statistical difference between control and mutant (SI Appendix, Figure S16).

**Figure 5.**
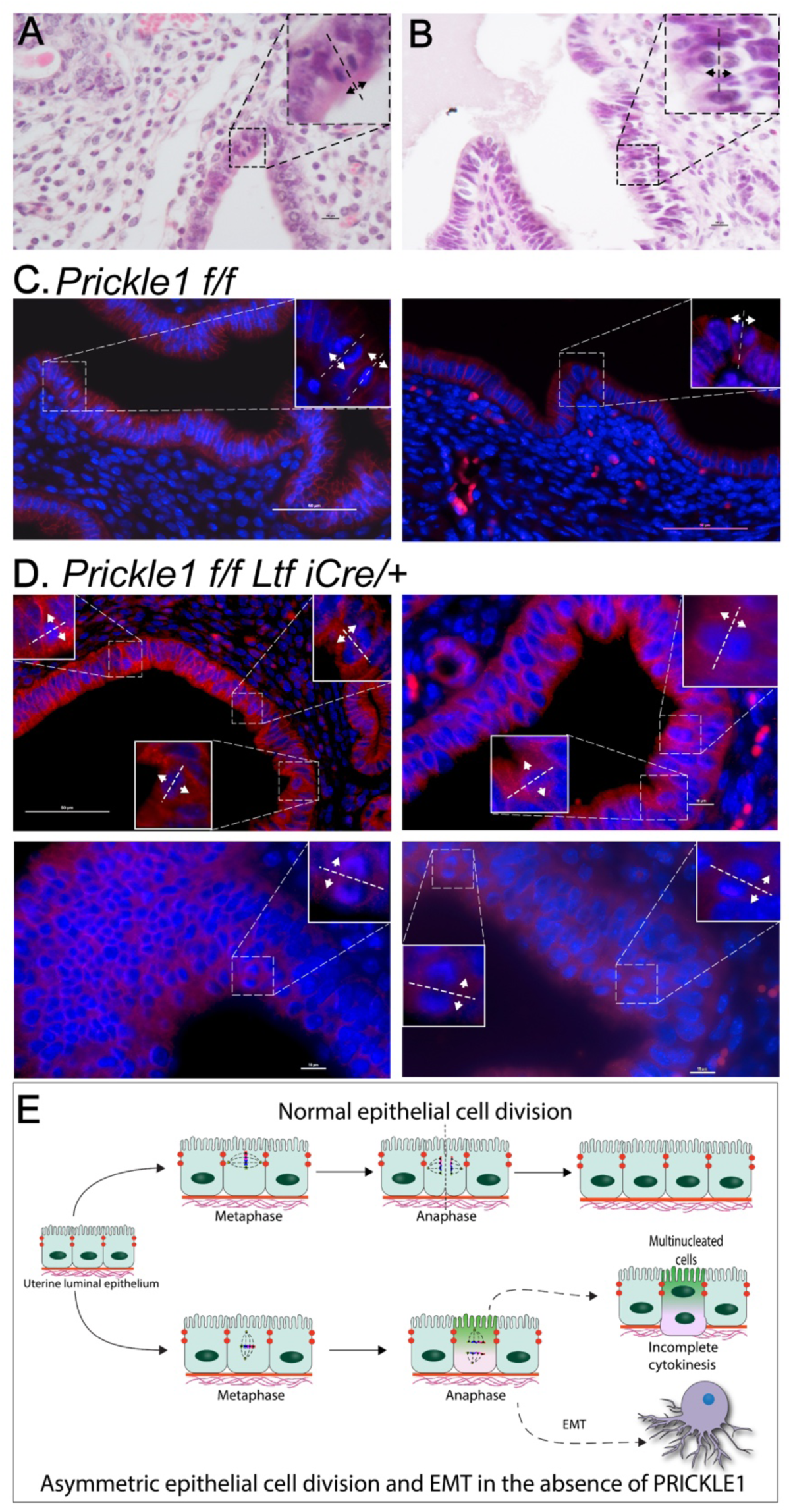
Loss of *Prickle1* under the *Ltf* promoter disrupts the polarity of cell division. (A,B) H&E stain of 6-month-old *Prickle1^f/f^ Ltf^+/icre^* cKO (B) compared to control (A) in diestrus. Callouts highlight dividing cells with altered orientation in the *Prickle1^f/f^ Ltf^+/icre^* cKO. (Scale bars, 10 μm). (C) Immunofluorescence stain of 6-month-old *Prickle1^f/f^* controls in diestrus for epithelial marker E-cadherin (red) and DAPI (blue). Callouts highlight dividing cells with normal orientation. (Scale bars, 50 μm). (D) Immunofluorescence stain of 6-month-old *Prickle1^f/f^ Ltf^+/icre^* cKO mice in diestrus for epithelial marker E-cadherin (red) and DAPI (blue). Callouts highlight dividing cells with altered orientation. (Scale bars, 50 μm). (E) Longitudinal view schematic depicting normal epithelial cell division and asymmetric epithelial cell division in the absence of PRICKLE1.

## Discussion

The mechanisms that regulate the unicellular layer architecture of endometrial luminal epithelium, timing, and cycle of the uterine receptive state for proper embryo implantation are poorly understood. In addition, although some recent advancements in understanding PCP-related functions within the uterus have been made (14), the specific role of PCP pathway in uterine epithelial morphogenesis and embryo implantation has previously been understudied. Here, we demonstrate the importance of PRICKLE1 in uterine endometrial epithelial architecture, embryo implantation, fertility, and overall uterine physiology using a novel *Prickle1* endometrial epithelial cKO mouse model.

We observed two distinct sets of phenotypes related to fertility in PRICKLE1 mutants, with a more severe mutant phenotype exhibiting no implantation sites and a less severe mutant with implantation sites similar to the control at GD 4.5 but with lower live pups at birth. Previous work with *lactoferrin-icre* has demonstrated known variability in the expression, leading to variability excision within mutants (25). The more severe mutants also displayed aberrant luminal folding, an open lumen, and altered AQP2 expression, indicating that *Prickle1^f/f^ Ltf^+/icre^*cKO mice may be unable to remove fluid from the lumen properly, thus preventing luminal closure and inhibiting successful embryo implantation. AQP2 is known to regulate luminal closure (27–29). Moreover, the increased expression of fibroblast growth factors, *Muc-1, Pgr,* and *Esr1* (SI Appendix, Figure S2 and S4) is consistent with results suggesting an impairment in the crosstalk between stroma and epithelium via Hand2 and progesterone (33). Decreased litter size even in mice that show normal implantation numbers at GD 4.5 may indicate this altered epithelial stromal crosstalk affects maintenance of pregnancy. The role of PRICKLE1 in epithelial-stromal communication and steroid hormone signaling needs further investigation.

Immunostaining of *Prickle1^f/f^Ltf^+/icre^*uteri for E-Cadherin and FOXA2 demonstrated glandular structure consistent with pregnancy at GD4.5 (Figure 2 C), even with an open lumen in the mutants that did not contain any positive embryo implantation sites. These results indicate that the inability of the lumen to fold and close properly, in addition to dysregulated cellular communication, may be responsible for the embryo implantation failure within these mice that otherwise should have been pregnant. Although the structure of the uterine glands of the mutant uteri appeared similar to the tubular and branched glands of control uteri, there were differences in the gland length that correlated with the open or closed luminal phenotype (Figure 2 C and D). The potential role of PRICKLE1 in regulating uterine gland morphogenesis needs further characterization.

Our scRNA sequencing analysis demonstrates how the loss of PRICKLE1 in the endometrial epithelium alters uterine gene expression and provides evidence for decreased number of cells with epithelial phenotype in the cKO with an increase in stromal cell numbers. Furthermore, RNA velocity analysis indicates a potential transition of epithelial cells into clusters with stromal-like gene expression. It has been well established that EMT is a contributor to several endometrial diseases, including endometriosis, adenomyosis, and endometrial cancers (52–54). Our data indicate a possible role for the loss of PRICKLE1 in EMT. In addition to the changes in epithelial and stromal cells in the cKO uteri, the scRNA data showed a significant increase in natural killer (NK) cells (AI Appendix, Figure S12 C). The over-abundance of NK cells in the cKO provides a potential connection to the fertility and implantation phenotype as it has been shown that uterine NK cell population dysregulation is connected to recurrent miscarriage, fertility, and uterine disorders such as endometriosis (55–57).

Spatial cues that organize a unicellular 2-dimensional sheet of epithelium in a plane orthogonal to the apical-basal polarity are regulated locally through various Wnt/ PCP signaling molecules (58). Additionally, symmetric (conservative) cell division in the epithelium is essential to maintain the apical-basal polarity and proper epithelial function, including implantation (4, 58). Our data indicating the presence of di-nucleated and multinucleated cells in the luminal epithelium, along with altered plane of cell division, indicate a crucial role for PRICKLE1 in maintaining conservative, symmetrical cell division in the epithelium and potentially in the maintenance of unicellular layer architecture. The presence of binucleated cells in the luminal epithelium may indicate incomplete cytokinesis or endoreduplication in cells (59). Proper cytokinesis requires the formation of a contractile ring at the site of E-cadherin expression (58). Therefore, increased basal E-cadherin expression and an altered mitotic plane of division support incomplete cytokinesis in our cKO. In this regard, regulation of expression and localization of E-cadherin by PRICKLE1 might hold the key for altered epithelial architecture and function. Furthermore, multinucleated spindle-shaped cells in the cKO epithelium may indicate that both defective cytokinesis as well as endoreduplication might be present (59). This observation, further strengthened by the increased number of doublets removed during scRNA data analysis in the cKO (SI Appendix, Table S16), is unique compared to the loss of other PCP genes such as *Vangl2, Ror2,* and *Wnt5a* (60, 61). Loss of these PCP genes has shown varying degrees of uterine epithelial subcellular changes as well as implantation defects. However, none of these genes have shown a dramatic loss of polarity of cell division or incomplete cytokinesis in the uterus. Moreover, the role of asymmetric cell division which gives rise to two daughter cells with distinct fates, in EMT has been reported (62). Whether the altered plane of cell division observed in the cKO endometrium has a direct role in the dramatic increase in EMT is unclear at this point.

## Materials and Methods

### Generation of *Prickle1* Conditional Knockout Mouse

Mouse embryonic stem cell clones harboring floxed *Prickle1* (*Prickle1tm1a(EUCOMM)Wtsi,* ES cell clones with conditional knockout potential, targeting vector HTGR03009_Z_6_E08) were acquired from EuCOMM. The ES cells were used to generate chimeric founder mice, which were mated to C57BL6 WT mice to create heterozygous *Prickle1*^f/+^ mice. These mice were crossed with FLPo (Gt(ROSA)26Sor^tm2(FLP*)Sor^) mice (JAX stock #009086) to remove FRT flanked bgal-neo sequences (63). These were bred together to obtain homozygous *Prickle1^f/f^*mice, which were then crossed with transgenic *lactoferrin-icre* mice (24). The mice were genotyped by PCR using primers for *Prickle1* (5’GGTTTCATGTGTTGAGACATTTC) (5’GTATTTCTGTGCCCTTTTTGTCGTCG) (5’TGAACTGATGGCGAGCTCAGACC) as well as primers for *Ltf^+/icre^* (5’AACTAGCACACCTGGTTGAGG) (5’CTTCTTGGGAGGCAGTGAAC) (5’CAGGTTTTGGTGCACAGTCA).

### Fertility and Implantation

Female *Prickle1^f/f^ Ltf^+/icre^* and *Prickle1^f/f^* mice were mated with C57BL6 WT males to induce pregnancy. The morning of finding the vaginal plug was considered day 0.5 of pregnancy. Litter size analysis was tracked via number of pups born following positive vaginal plug formation at days 17-21. Implantation was assessed by sacrificing pregnant dams on day 3.5 or 4.5 following 100 μL intravenous injection of 1% Chicago Sky Blue 6B dye (Millepore Sigma, Cat# C8679) solution and counting the number of distinct blue bands present within the uterus.

### Whole mount immunofluorescence, confocal imaging and image analysis

Whole-mount immunofluorescence was performed as previously described (13). GD 4.5 *Prickle1^f/f^* and *Prickle1^f/f^ Ltf^+/icre^ uteri* were dissected from mice and fixed in DMSO:methanol (1:4). Subsequently, they were rehydrated in a methanol:PBST (1% Triton X-100 in PBS) (1:1) solution for 15 minutes, followed by a PBST wash for 15 minutes. Uteri were then incubated in a blocking solution (2% powdered milk in PBST) for 1 hour at room temperature. They were incubated with primary antibodies diluted in blocking solution for nine nights at 4°C. Primary antibodies included rabbit anti-FOXA2 (Abcam, ab108422; 1:300) and rat anti-CDH1 (M108, Takara Biosciences, 1:500). Uteri were washed once for 15 minutes with 1% PBST followed by four additional PBST washes for 45 minutes each at room temperature. Uteri were then incubated with 1:500 Alexa Fluor 555 donkey anti-rabbit IgG (Invitrogen, A31572), Alexa Fluor 633 goat anti-rat IgG (Invitrogen, A21094) and Hoechst (Sigma Aldrich, B2261) at 4°C for three nights. Samples were washed once for 15 min and three more times for 45 minutes each with 1% PBST, and then stepwise dehydrated into 100% methanol. Uteri were incubated overnight at 4°C in a 3% H_2_O_2_ solution prepared in methanol. Next day, the samples were washed twice for 15 minutes each and a final 1 hour wash in 100% methanol at room temperature, and then cleared in BABB (1:2, benzyl alcohol:benzyl benzoate) (Sigma-Aldrich, 108006, B6630) overnight.

#### Confocal Imaging

Whole tissue immunofluorescence samples were imaged using a Leica TCS SP8 X Confocal Laser Scanning Microscope System with white-light laser, 10X air objective and 20X BABB objective. Using the tile scan function with Z stacks 7μm apart (10X), images covering the entire length and thickness of the uterine horn were acquired. Higher resolution images of implantation sites and regions flanking the implantation site were acquired using a 20X BABB objective. The tile scan function was used and Z stacks 5μm apart were acquired.

#### Image Analysis

Image analysis was performed using commercial software Imaris v9.2.1 (Bitplane). The confocal image (.LIF) files were imported into the Surpass mode of Imaris. To derive the lumen-only signal, the FOXA2 signal of glands was subtracted from the epithelial CDH1 signal using the channel arithmetics Matlab-based function (26). 3D renderings were created using automated and manual mode in Surface function for lumen-only and FOXA2 or CDH1 (glands only) signal. Individual glands were then isolated and presented in a comprehensive gallery for visualization using the Imaris Vantage function. Gland length was determined using Bounding Box OOC function in Imaris that measures the shortest straight-line distance from its point of connection to the uterine lumen to the furthest tip. To compare gland length measurements between *Prickle1*^f/f^ mice and *Prickle1*^f/f^ *Ltf*^+/icre^ cKO mice, Kruskal-Wallis test with Dunn’s multiple comparisons was performed using R Statistical Software. p-value <0.05 was considered significant indicating differences between comparisons.

### Histology and Staining

Uterine tissues were fixed in 4% paraformaldehyde and processed for paraffin embedding. Tissue sections were deparaffinized in xylene and rehydrated using a series of ethanol and then stained with Hematoxylin and Eosin (H&E) or immunostained. For immunostaining, antigen retrieval was done using antigen retrieval unmasking solution (Vector Labs, Cat# H-3301) prior to staining.

For immunofluorescence staining, rehydrated tissue sections were stained as previously described (64) using primary antibodies AQP2 (Novus Biologicals, Cat# NB110-74682SS, 1:100), E-CAD (Cell Signaling, Cat# 24E10, 1:100), ZEB1 (Cell Signaling, Cat# 70512, 1:400), TWIST1 (Cell Signaling, Cat# 90445, 1:100), and SNAI2 (Cell Signaling, Cat# 9585, 1:100). Secondary antibodies were obtained from Jackson Immuno Research Laboratories for Alexa Fluor 488 and 594 (Cat# 711-545-152 and 711-585-152, 1:200).

For immunohistochemistry staining, rehydrated samples were prepared using an ABC Universal PLUS Peroxidsae kit (Vector Labs, Cat# PK-8200) and then with primary antibodies KRT7 (ThermoFisher Scientific, Cat# pA5-82291, 1:500), SOX9 (Cell Signaling, Cat# 82630, 1:100), and Vimentin (Cell Signaling, Cat# D21H3, 1:100).

For TUNEL Apoptosis staining, rehydrated samples were prepared using a One-Step TUNEL In Situ Apoptosis kit (ELabscience, Cat# E-CK-A320) and then imaged.

### RNA Isolation and qRT-PCR Analysis

Total RNA was isolated from uterine tissue samples stored in RNAlater (Invitrogen, Cat# AM7020) and then biopulvirized and placed into TRIzol Reagent (Invitrogen, Cat# 15596026). Following TRIzol, a series of chloroform, isopropanol, and ethanol washes were used to isolate the RNA. Following quantification using Nanodrop spectrophotometer, aliquots of RNA were reverse transcribed using High-Capacity cDNA Reverse Transcription Kit (Applied Biosystems, 4368814). TaqMan assays for *Fgf1* (IDT, Mm.PT.56a.41158563)*, Fgf2* (IDT, Mm.PT.56a.5129235)*, Fgf7* (IDT, Mm.PT.56a.28834565)*, Fgf9* (IDT, Mm.PT.56a.5456225)*, Fgf10* (IDT, Mm.PT.58.11905869), and *Fgf18* (IDT, Mm.PT.58.14021387), and *Muc-1* (IDT, Mm.PT.58.15865847) were used to quantify gene expression differences utilizing the delta delta C(T) method with housekeeping genes Rn18s (ThermoFisher Scientific, Mm03928990_G1).

### Hormone Assessment

Samples of serum from *Prickle1^f/f^ Ltf^+/icre^* (n=4) and *Prickle1^f/f^* (n=4) mice aged 4 months were assayed by ELISA at the University of Virginia Center for Research in Reproduction, Ligan Assay and Analysis Core (Charlotteville, VA).

### Single-cell RNA Sequencing

Whole uteri of *Prickle1^f/f^ Ltf^+/icre^* (n=3) and *Prickle1^f/f^* (n=3) mice aged 6 months were isolated immediately following euthanasia using IACUCC approved methods and washed in cold 1X PBS. Uteri were digested as previously described (41), and samples were pooled following digestion. The KUMC genomics core processed and sequenced the cells as previously described (41). The *Prickle1^f/f^ Ltf^+/icre^* sample had 5805 cells with 70202 mean reads and 2078 median genes per cell while the *Prickle1^f/f^* sample had 5426 cells with 43202 mean reads and 1098 median genes per cell. Both samples had high sequence saturation levels (> 60%) and the mapping rate of reads to the mouse genome (mm10) was > 90% for both samples.

### Single-cell Data Analysis

scRNA-sequencing libraries were generated using the 10x Chromium Single Cell 3’ v3 chemistry (10x Genomics) and sequenced in an Illumina NovaSeq 6000 sequencing machine. The raw sequence reads were processed using the 10x Genomics Cellranger pipeline (v 6.1.1) to obtain UMI feature-barcode count matrices. Doublets were removed using DoubletFinder (65) and cells filtered to contain only those cells with at least 500 UMIs and over 250 detected genes with a genes per UMI ratio greater than 0.8 and a mitochondrial gene ratio less than 20%. The resulting single cell data was analyzed using the R software Seurat (66) (v4) as previously described (67). The analysis was performed at a 0.4 cluster resolution reasoned using the Clustree software (68) giving 12 stable clusters. The cluster cell types were identified using the SingleR software (39) and our expert curation as previously described (41). Annotations were based on the two reference datasets, ImmGenData from the Immunological Genome Project (ImmGen) (69) and MouseRNAseqData (70). The dynamic progression of transcription in the single cell data were analyzed using the Velocyto package (49).

## Acknowledgments and Funding Sources

V.M.C. was supported by grants from the NIH: P20 RR016475, R01 HD094373, R01 HD076450, R01 HD105714-01. R.A. was supported by grants from NIH: R01 HD109152. E.R.R. was supported by the Madison and Lila Self Graduate Fellowship. A.V.B. was supported by the Michigan State University, Department of Obstetrics, Gynecology, and Reproductive Biology, Walstrom Family Endowed Women’s Health Research Fund.

The authors acknowledge the Transgenic and Gene-Targeting Institutional Facility for help with the transgenic mice, the Genomics Core (Kansas Intellectual and Developmental Disability Research Center (KIDDRC) (NIH U54 HD090216), the Centers of Biomedical Research Excellence (P30 GM122731-03), and the NIH S10 High-End Instrumentation Grant (NIH grants S10 OD021743)) for help with the scRNA sequencing, and the KIDDRC (NIH U54 HD 090216) for help with the imaging at the University of Kansas Medical Center, Kansas City, KS, 66160. The authors also thank the University of Virginia Center for Reproduction Ligand Assay and Analysis core, supported by NIH/NCTRI P50-HD28934.

## Author Contributions

V.M.C and R.A designed research; E.R.R., A.V.B., S. Ganeshkumar, S. Gunewardena, and R.A. performed research; E.R.R., A.V.B, S. Gunewardena, R.A., and V.M.C. analyzed data; E.R.R., S. Gunewardena, A.V.B., R.A., and V.M.C wrote and edited the manuscript.

## Data Availability Statement

Upon acceptance of the manuscript for publication, the authors agree to publicly release all data underlying the study. This includes single-cell RNA sequencing data submitted to NCBI GEO. The codes used for data analysis are included in the materials and methods.

