## Supplemental Information Appendix 2 for "Loss of PRICKLE1 leads to abnormal endometrial epithelial architecture, decreased embryo implantation, and reduced fertility in mice"

### This PDF file includes:

Appendix 2 Figures 1A-L

### Other supporting materials for this manuscript include the following:

SI Appendix

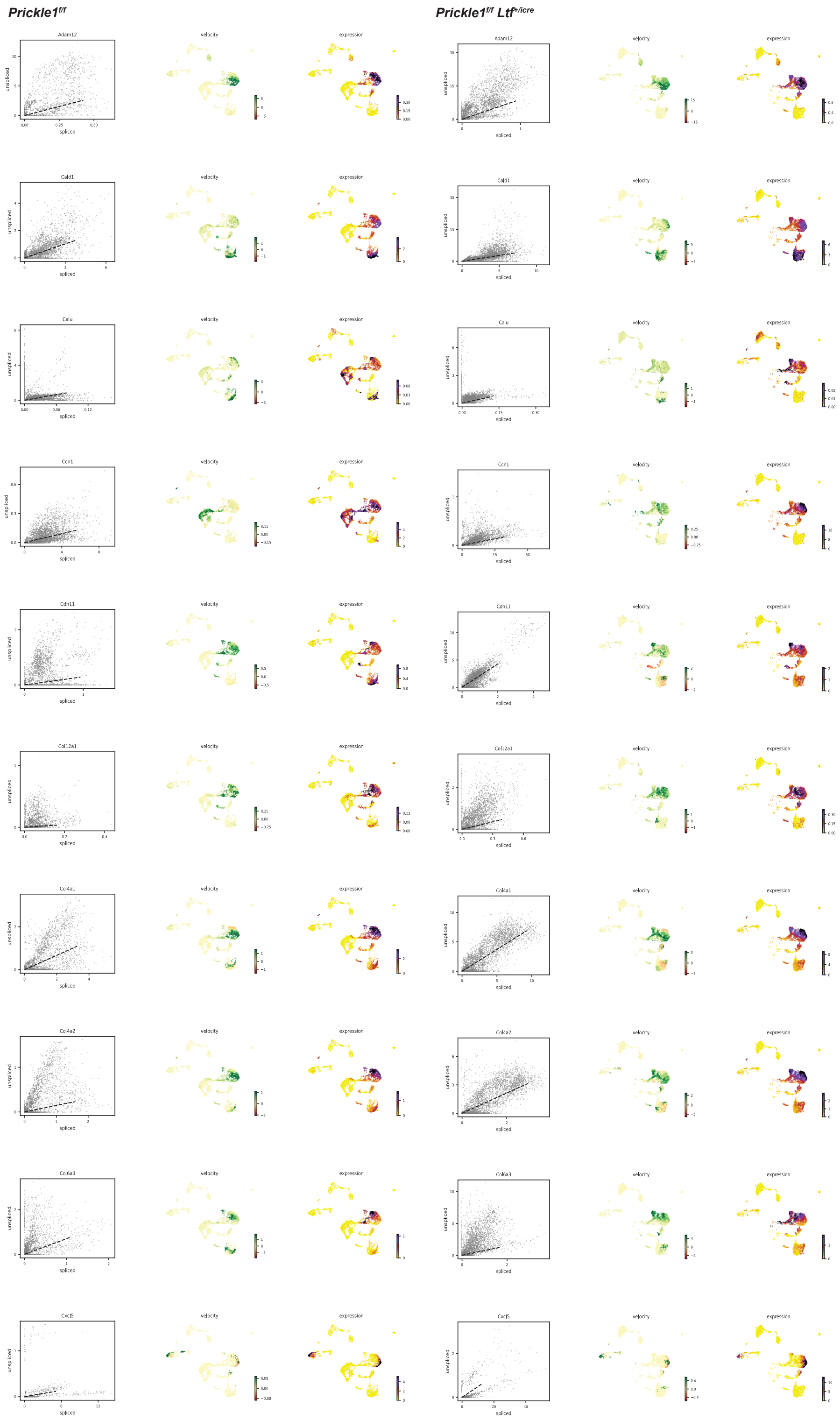


**Appendix 2 Figure 1A.** <u>Loss of *Prickle1* under the *Ltf* promoter results in high velocity of EMT hallmark genes in stromal cells (clusters 1 and 2).</u> Velocity dynamics of 120 hallmark EMT genes showing the ratio of spliced and unspliced RNA transcripts overall, velocity RNA analysis, and expression levels in each cluster for control and *Prickle1*$^{f/f}$ *Ltf*$^{+/icre}$ mice.

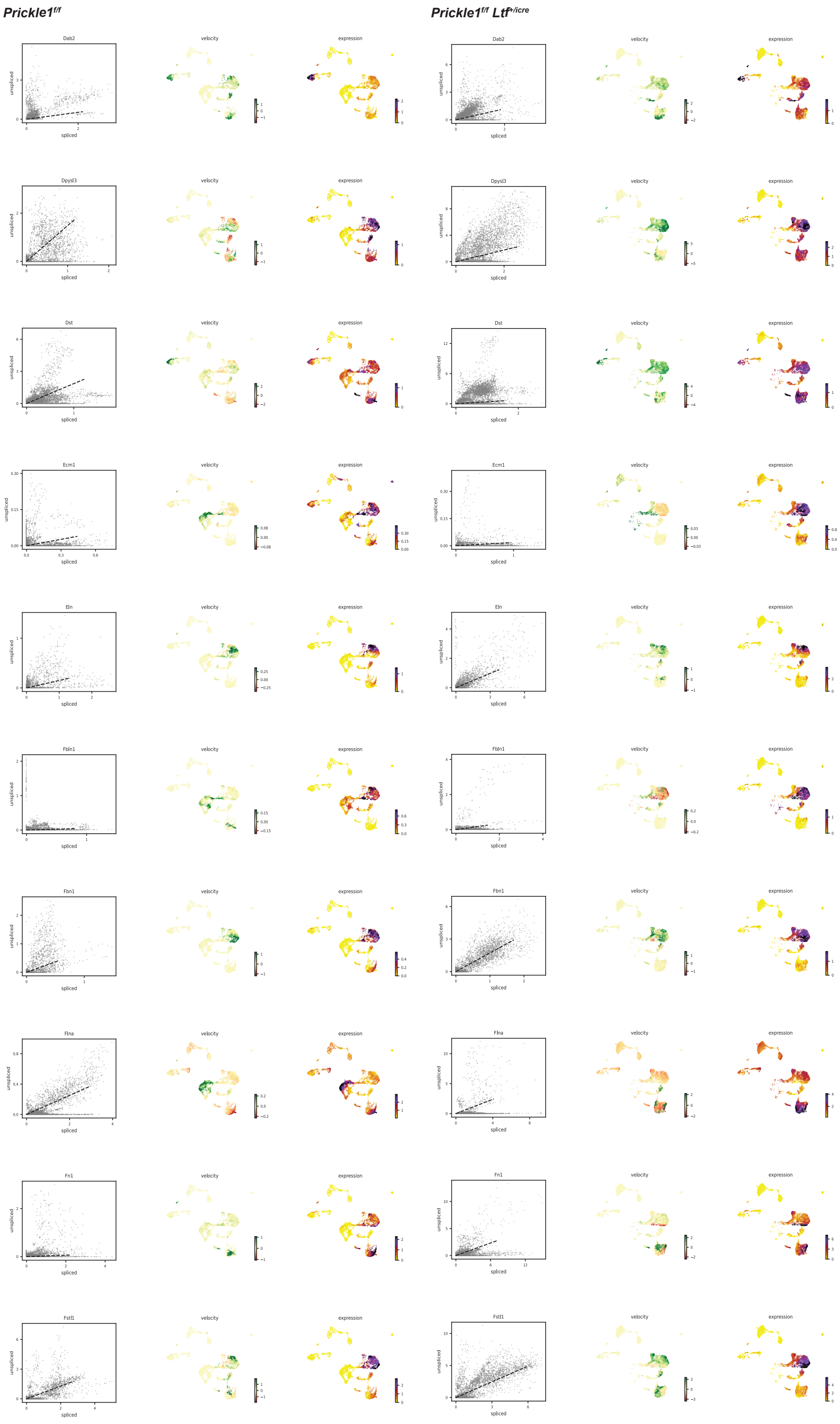


**Appendix 2 Figure 1B.** <u>Loss of *Prickle1* under the *Ltf* promoter results in high velocity of EMT hallmark genes in stromal cells (clusters 1 and 2).</u> Velocity dynamics of 120 hallmark EMT genes showing the ratio of spliced and unspliced RNA transcripts overall, velocity RNA analysis, and expression levels in each cluster for control and *Prickle1*$^{f/f}$ *Ltf*$^{+/icre}$ mice.

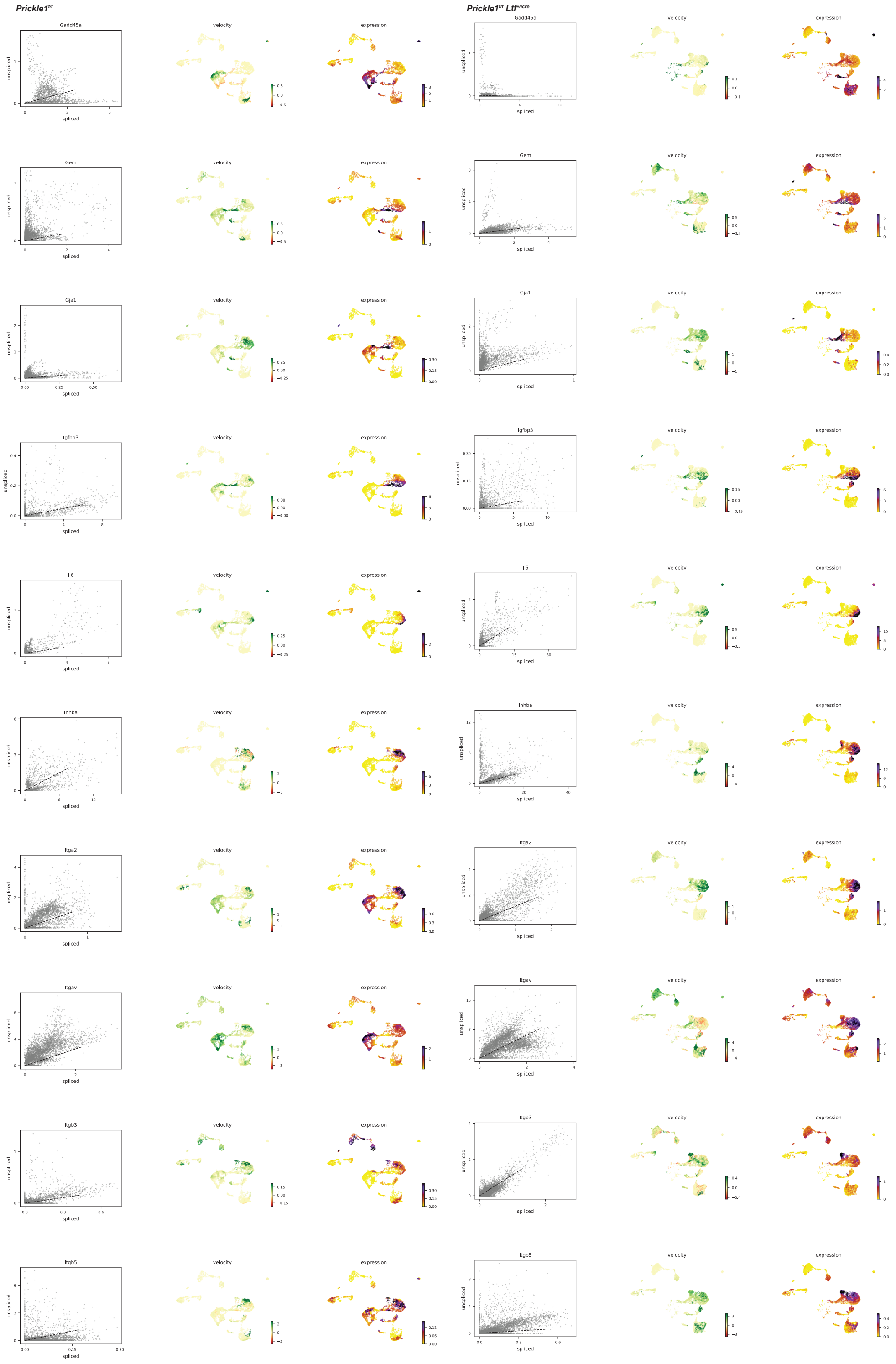


**Appendix 2 Figure 1C.** Loss of *Prickle1* under the *Ltf* promoter results in high velocity of EMT hallmark genes in stromal cells (clusters 1 and 2). Velocity dynamics of 120 hallmark EMT genes showing the ratio of spliced and unspliced RNA transcripts overall, velocity RNA analysis, and expression levels in each cluster for control and *Prickle1*$^{f/f}$ *Ltf*$^{+/icre}$ mice.

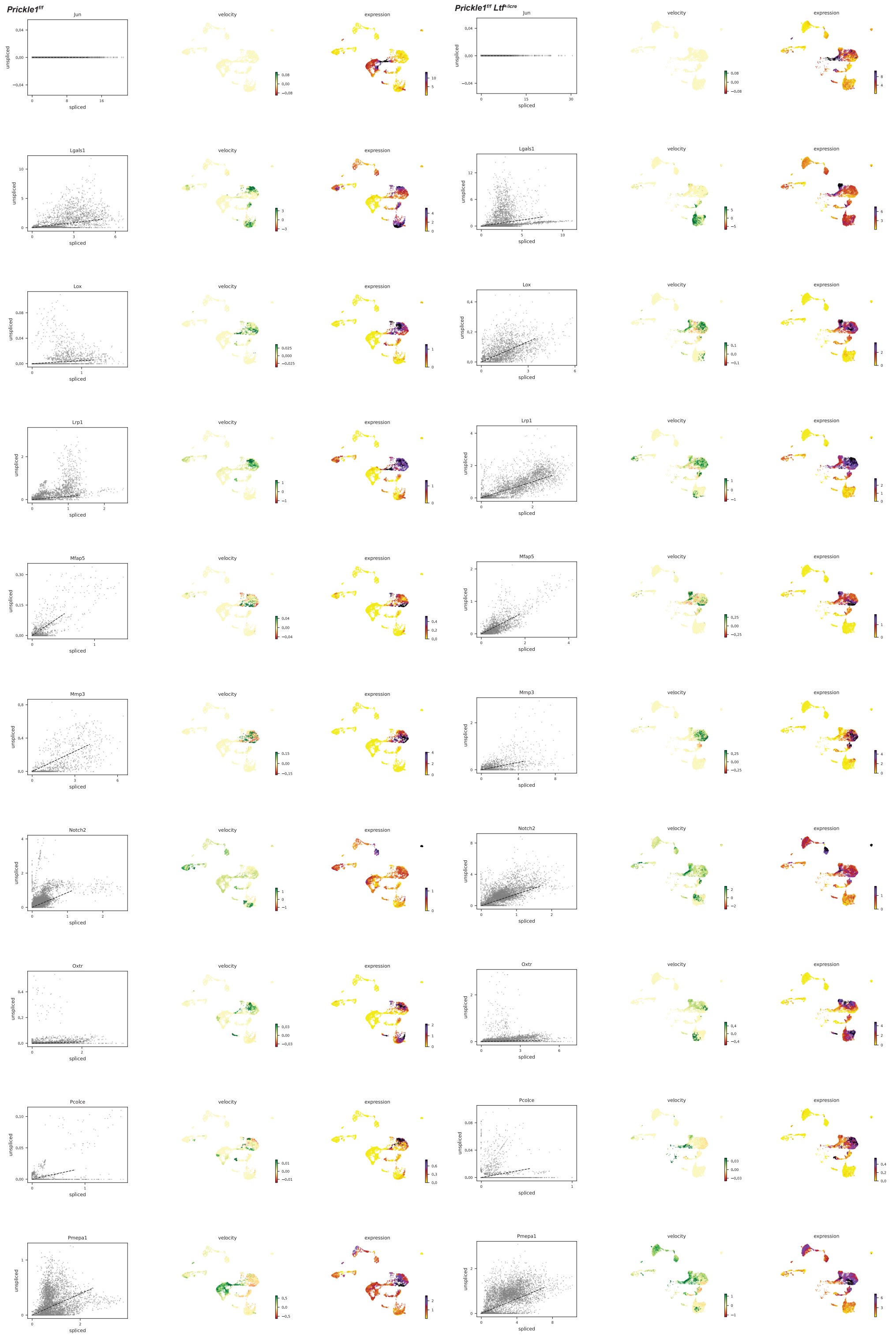


**Appendix 2 Figure 1D.** Loss of *Prickle1* under the *Ltf* promoter results in high velocity of EMT hallmark genes in stromal cells (clusters 1 and 2). Velocity dynamics of 120 hallmark EMT genes showing the ratio of spliced and unspliced RNA transcripts overall, velocity RNA analysis, and expression levels in each cluster for control and *Prickle1*$^{f/f}$ *Ltf*$^{+/icre}$ mice.

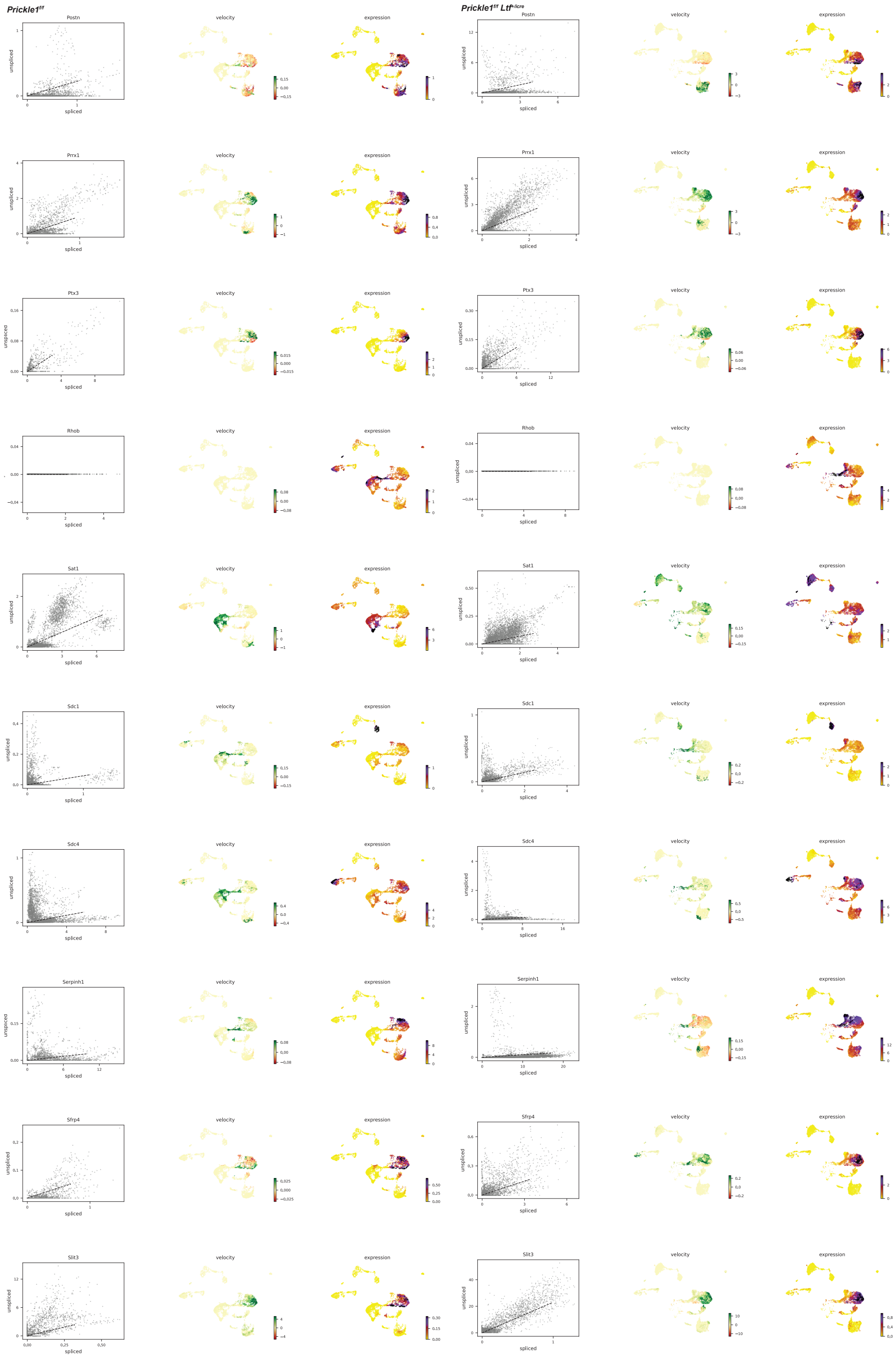


**Appendix 2 Figure 1E.** <u>Loss of *Prickle1* under the *Ltf* promoter results in high velocity of EMT hallmark genes in stromal cells (clusters 1 and 2).</u> Velocity dynamics of 120 hallmark EMT genes showing the ratio of spliced and unspliced RNA transcripts overall, velocity RNA analysis, and expression levels in each cluster for control and *Prickle1*$^{f/f}$ *Ltf*$^{+/icre}$ mice.

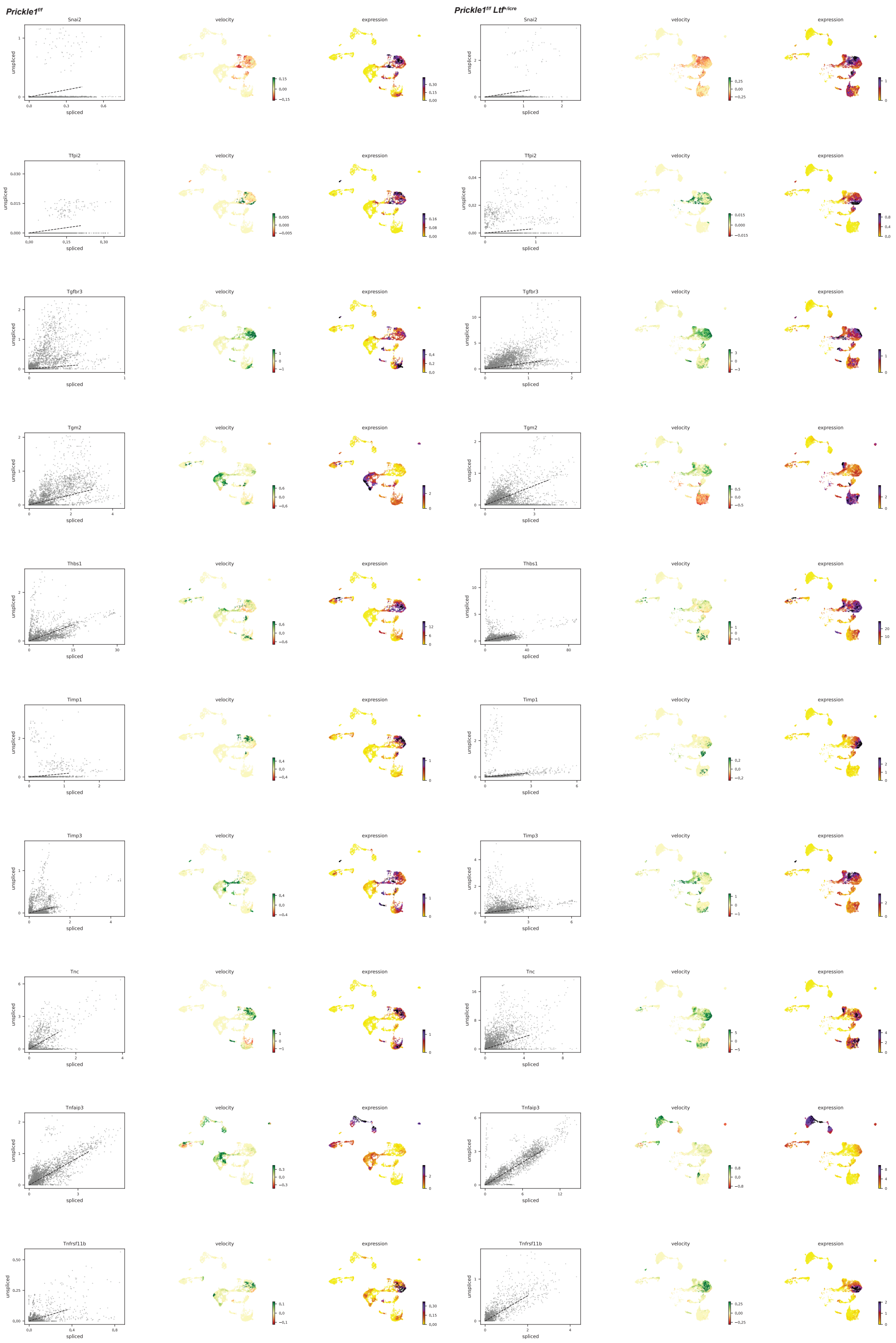


**Appendix 2 Figure 1F.** Loss of *Prickle1* under the *Ltf* promoter results in high velocity of EMT hallmark genes in stromal cells (clusters 1 and 2). Velocity dynamics of 120 hallmark EMT genes showing the ratio of spliced and unspliced RNA transcripts overall, velocity RNA analysis, and expression levels in each cluster for control and *Prickle1*$^{f/f}$ *Ltf*$^{+/icre}$ mice.

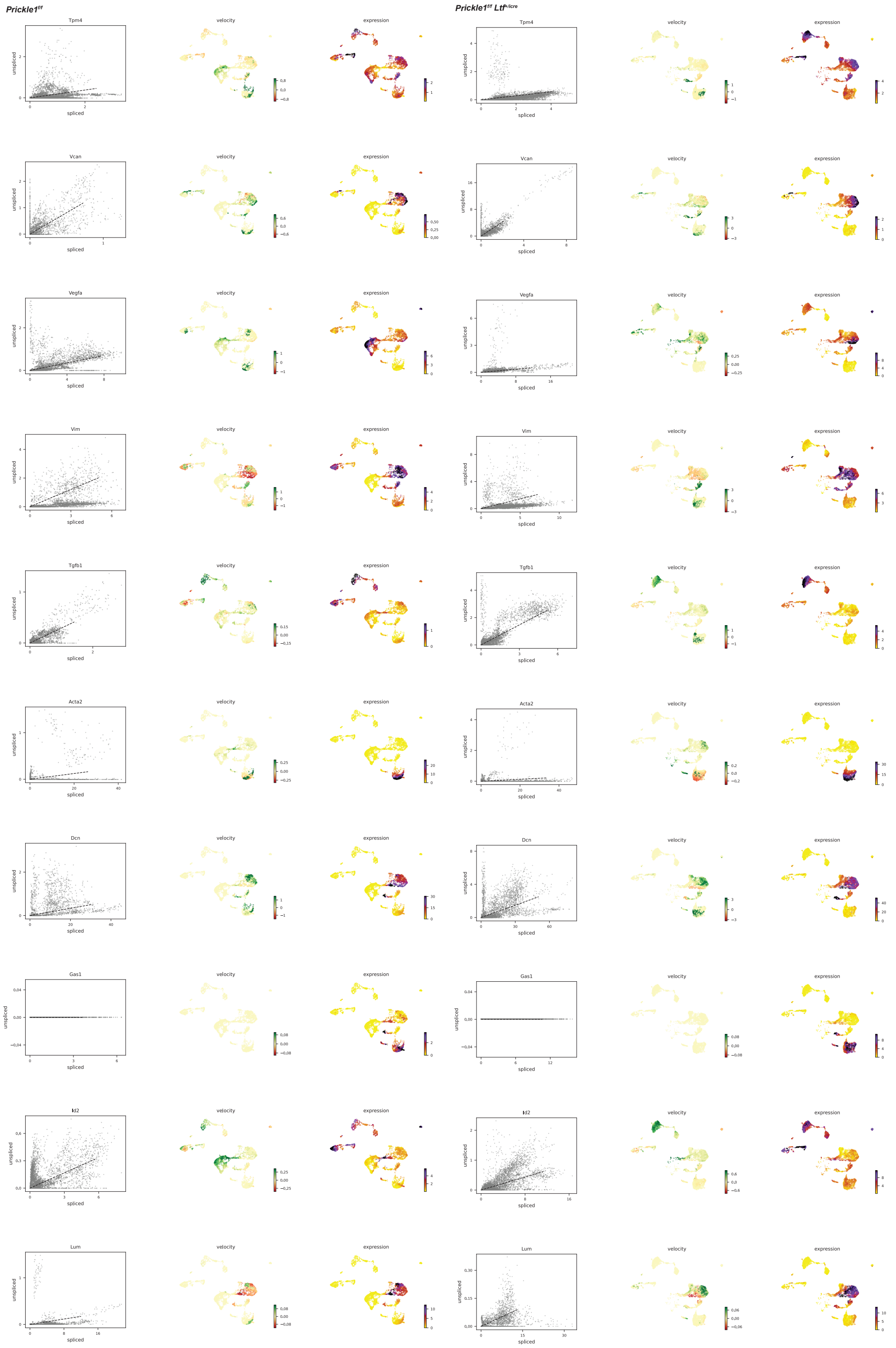

**Appendix 2 Figure 1G.** Loss of *Prickle1* under the *Ltf* promoter results in high velocity of EMT hallmark genes in stromal cells (clusters 1 and 2). Velocity dynamics of 120 hallmark EMT genes showing the ratio of spliced and unspliced RNA transcripts overall, velocity RNA analysis, and expression levels in each cluster for control and *Prickle1*$^{f/f}$ *Ltf*$^{+/icre}$ mice.

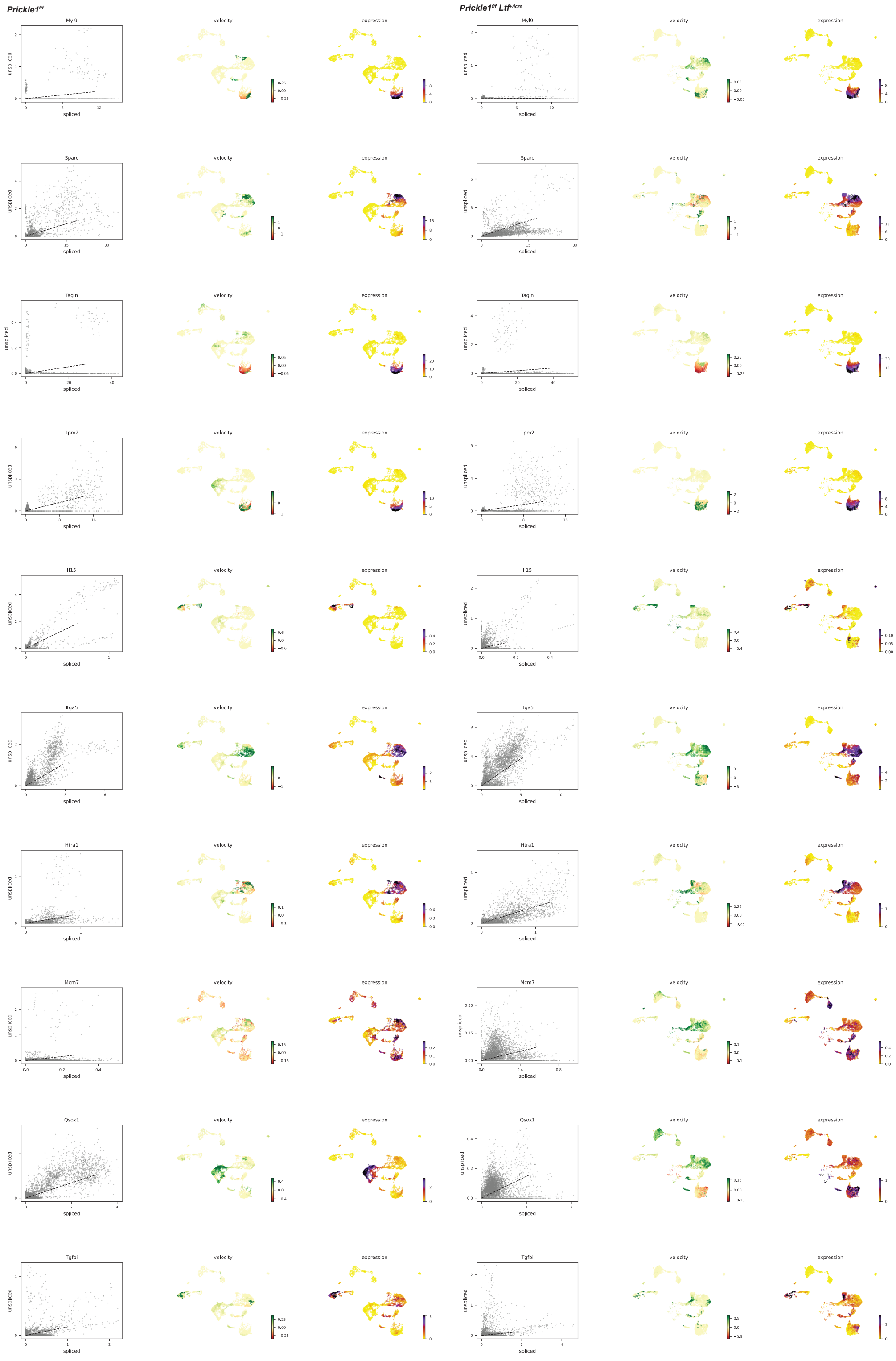


**Appendix 2 Figure 1H.** Loss of *Prickle1* under the *Ltf* promoter results in high velocity of EMT hallmark genes in stromal cells (clusters 1 and 2). Velocity dynamics of 120 hallmark EMT genes showing the ratio of spliced and unspliced RNA transcripts overall, velocity RNA analysis, and expression levels in each cluster for control and *Prickle1*$^{f/f}$ *Ltf*$^{+/icre}$ mice.

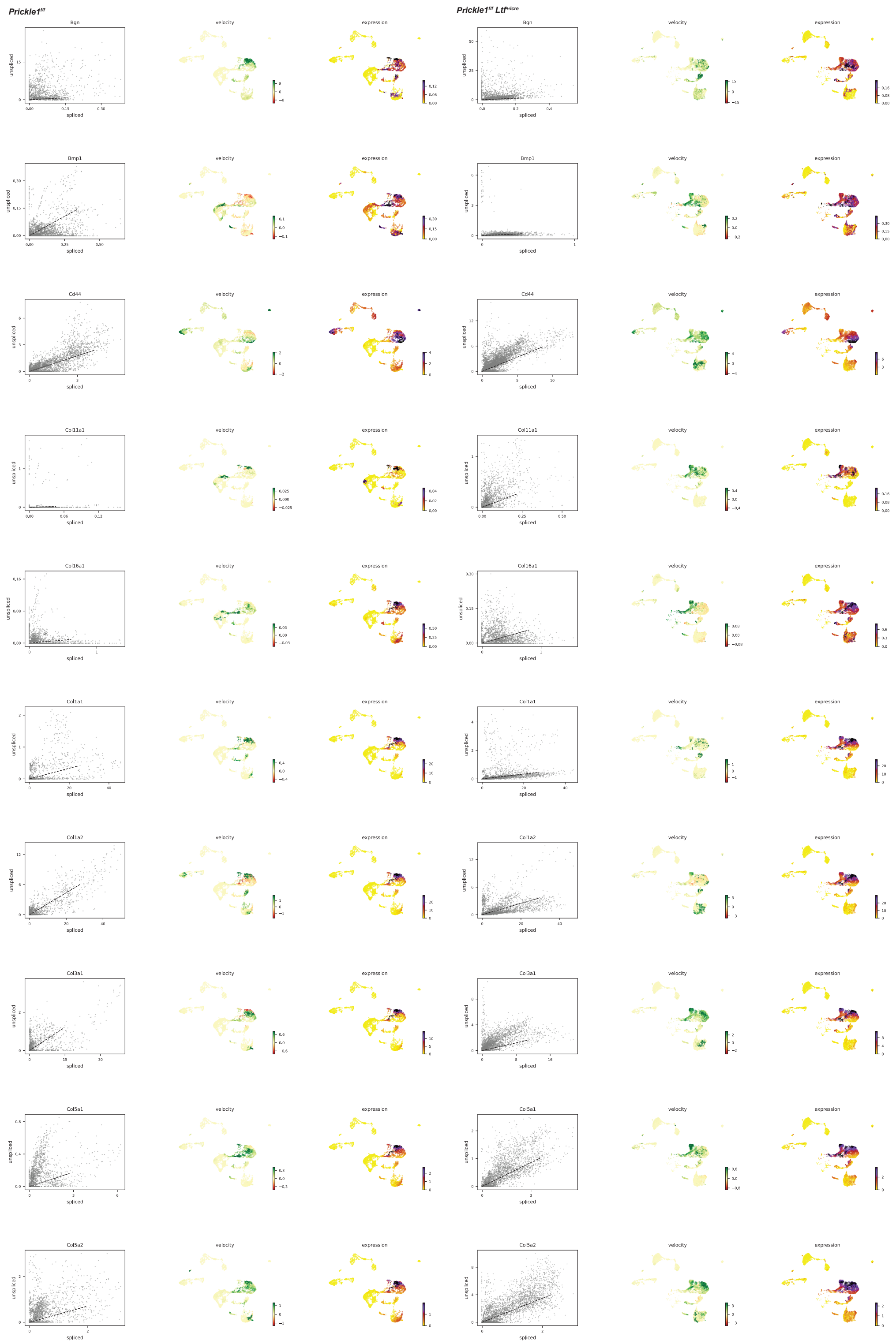


**Appendix 2 Figure 1I.** Loss of *Prickle1* under the *Ltf* promoter results in high velocity of EMT hallmark genes in stromal cells (clusters 1 and 2). Velocity dynamics of 120 hallmark EMT genes showing the ratio of spliced and unspliced RNA transcripts overall, velocity RNA analysis, and expression levels in each cluster for control and *Prickle1*$^{f/f}$ *Ltf*$^{+/icre}$ mice.

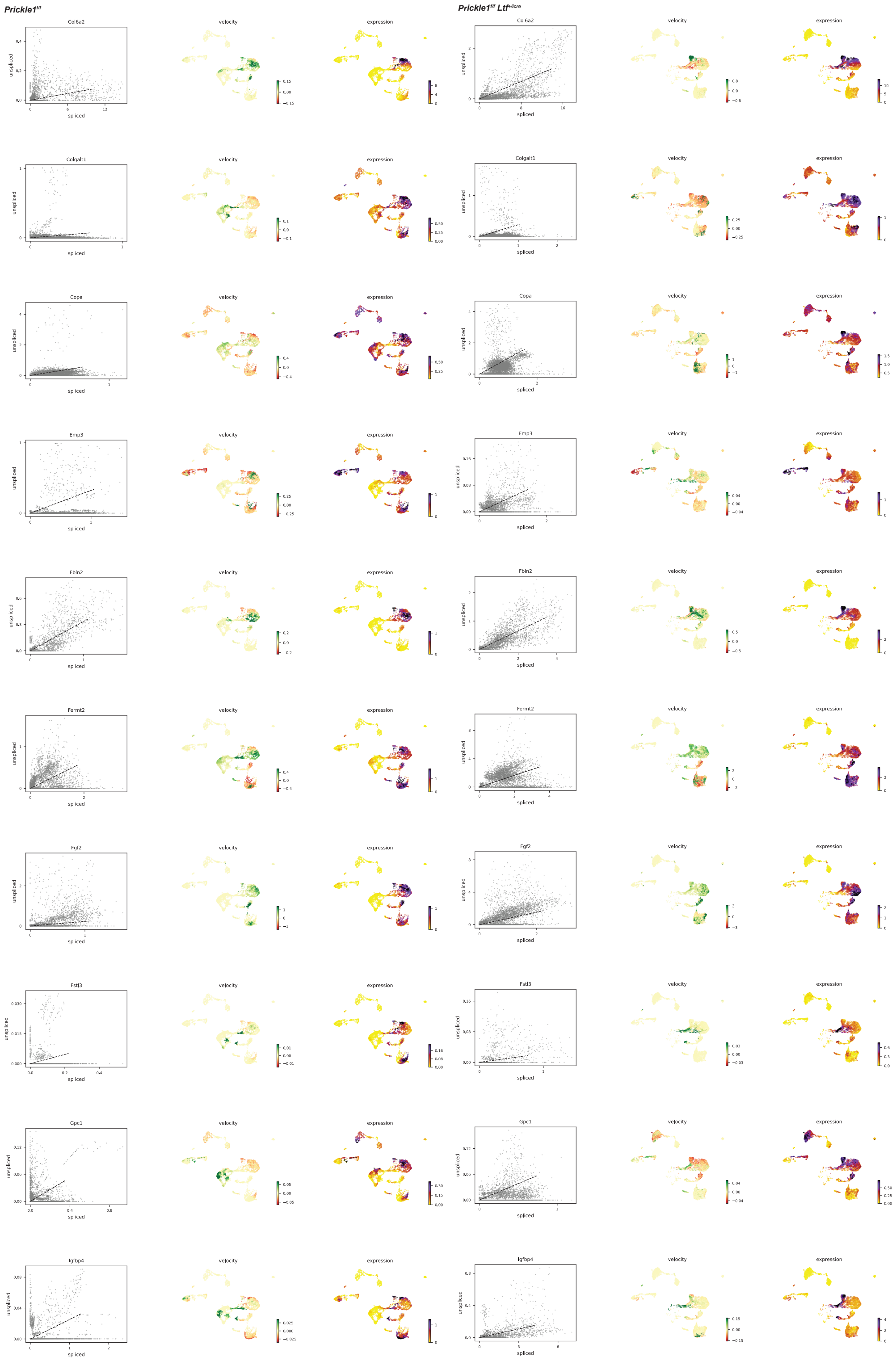


**Appendix 2 Figure 1J.** Loss of *Prickle1* under the *Ltf* promoter results in high velocity of EMT hallmark genes in stromal cells (clusters 1 and 2). Velocity dynamics of 120 hallmark EMT genes showing the ratio of spliced and unspliced RNA transcripts overall, velocity RNA analysis, and expression levels in each cluster for control and *Prickle1*$^{f/f}$ *Ltf*$^{+/icre}$ mice.

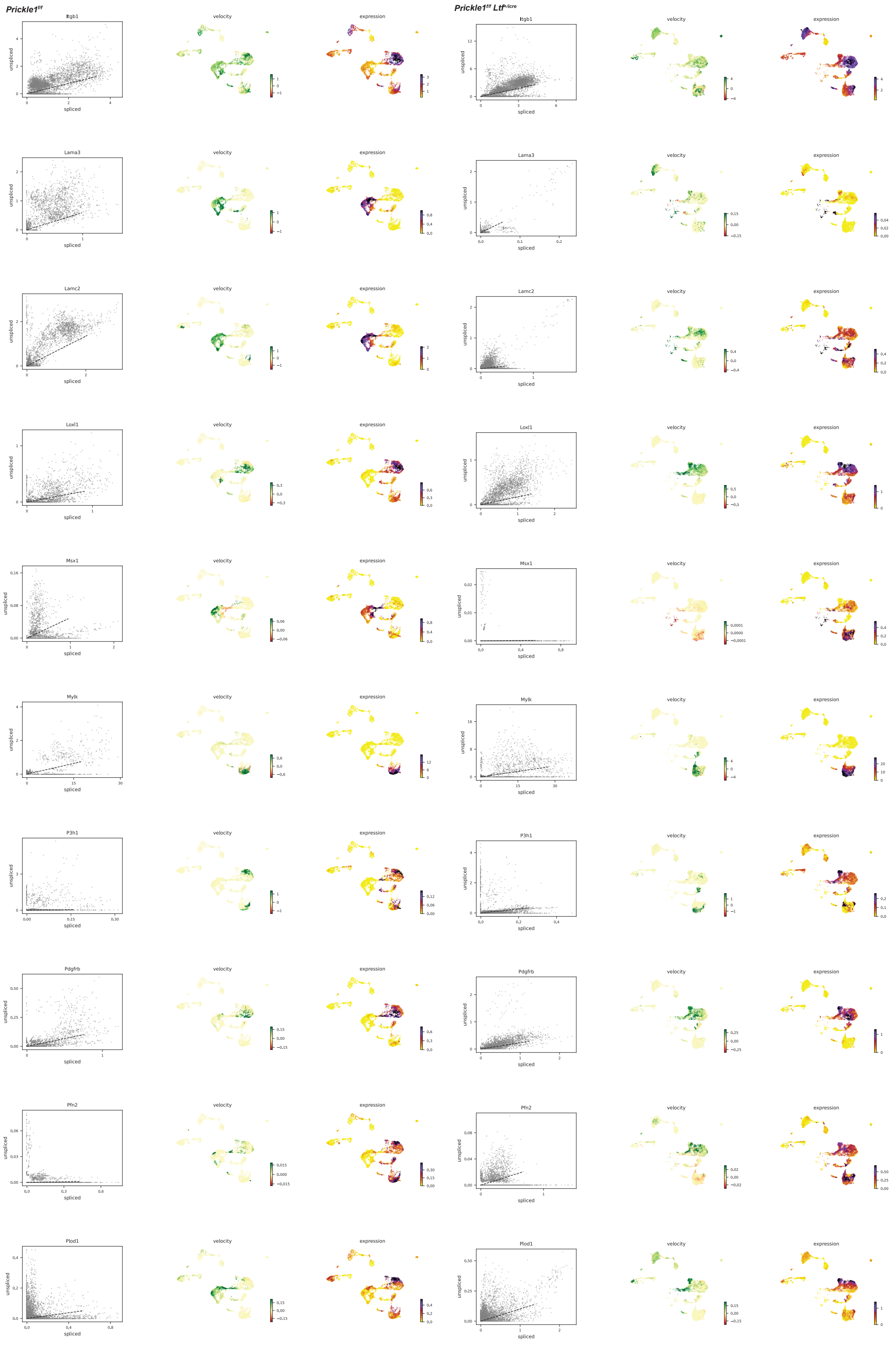


**Appendix 2 Figure 1K.** Loss of *Prickle1* under the *Ltf* promoter results in high velocity of EMT hallmark genes in stromal cells (clusters 1 and 2). Velocity dynamics of 120 hallmark EMT genes showing the ratio of spliced and unspliced RNA transcripts overall, velocity RNA analysis, and expression levels in each cluster for control and *Prickle1*$^{f/f}$ *Ltf*$^{+/icre}$ mice.

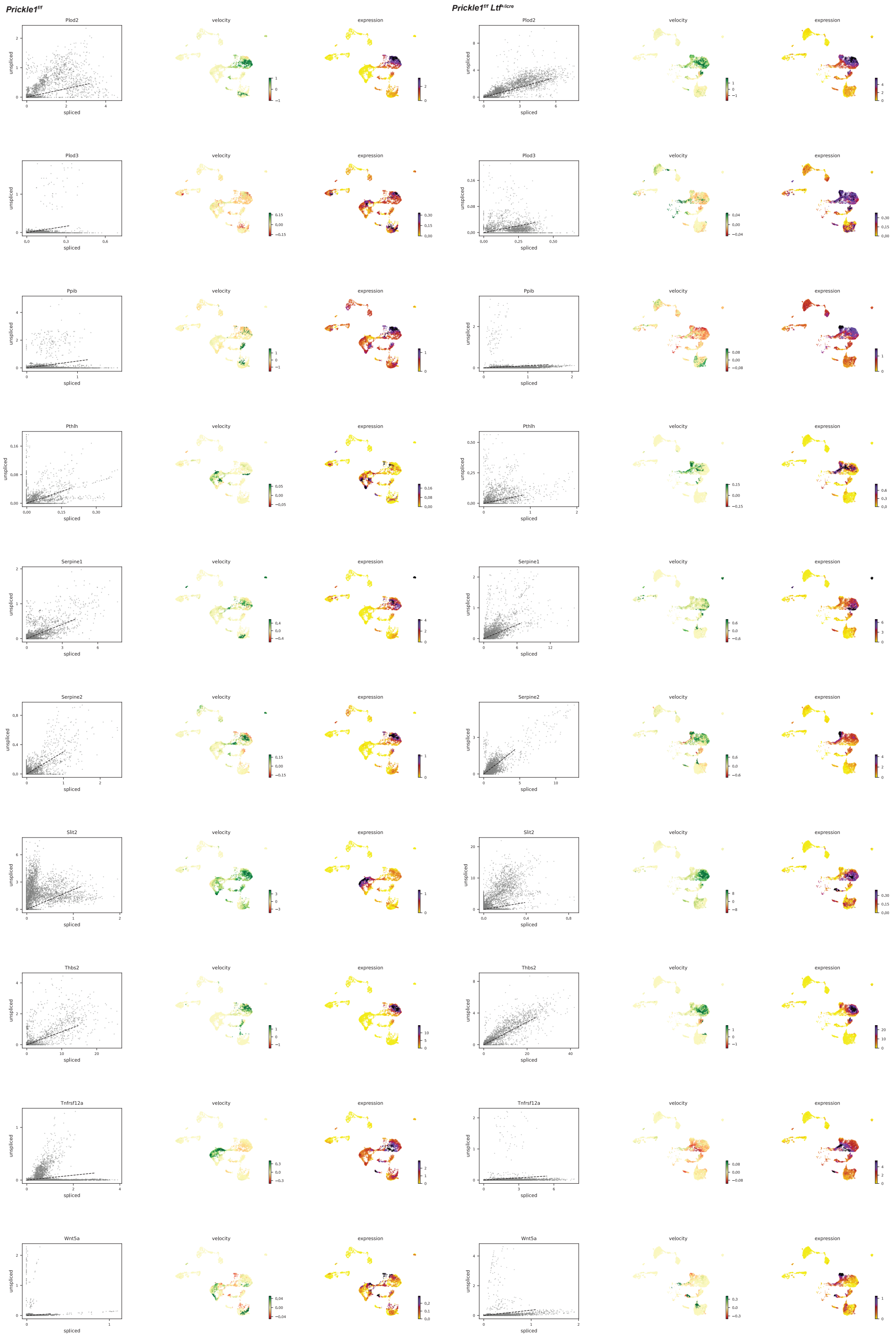


**Appendix 2 Figure 1L.** Loss of *Prickle1* under the *Ltf* promoter results in high velocity of EMT hallmark genes in stromal cells (clusters 1 and 2). Velocity dynamics of 120 hallmark EMT genes showing the ratio of spliced and unspliced RNA transcripts overall, velocity RNA analysis, and expression levels in each cluster for control and *Prickle1*$^{f/f}$ *Ltf*$^{+/icre}$ mice.
