## Supplemental Information Appendix for "Loss of PRICKLE1 leads to abnormal endometrial epithelial architecture, decreased embryo implantation, and reduced fertility in mice"

**This PDF file includes:**

Figures S1 to S16
Tables S1 to S17

**Other supporting materials for this manuscript include the following:**

SI Appendix 2

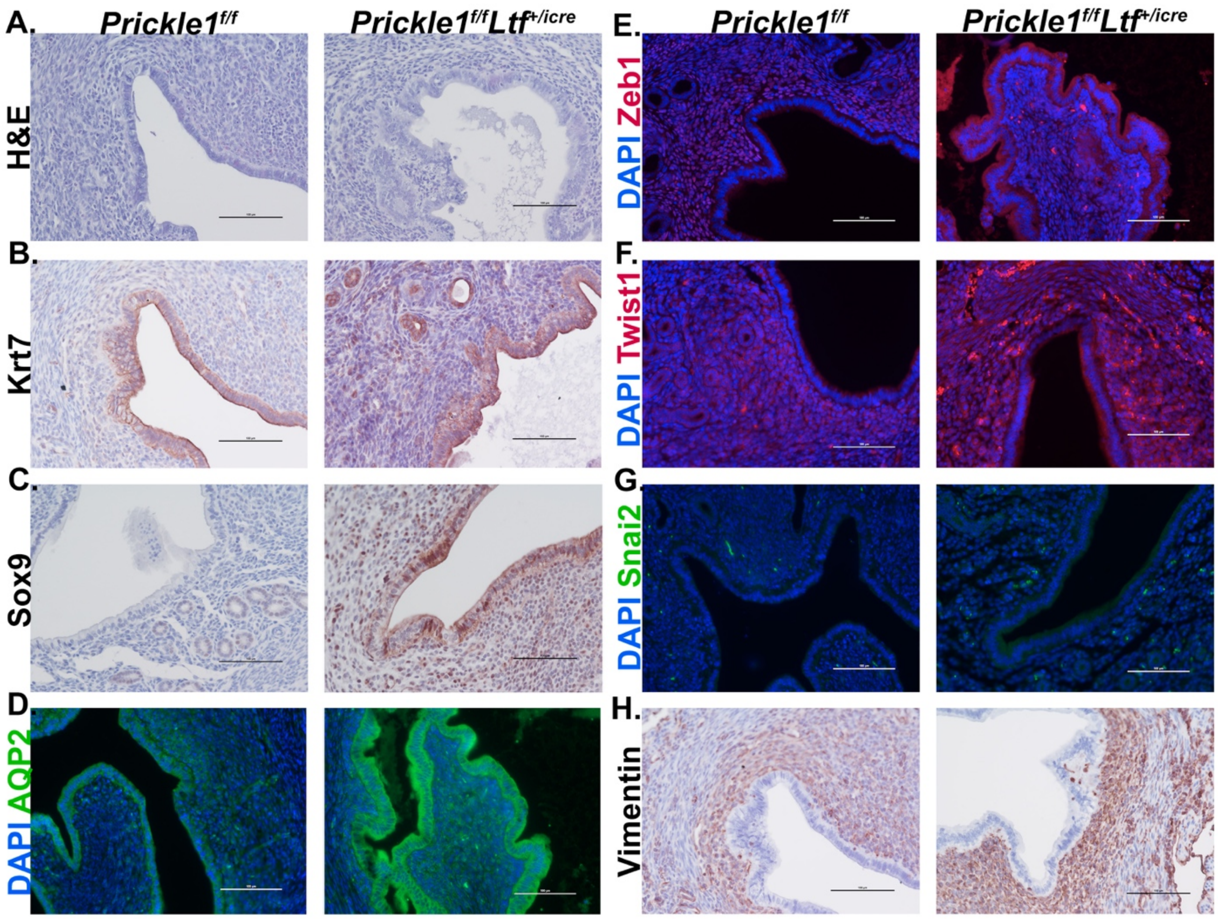


**Supplemental Figure 1.** Loss of *Prickle1* under the *Ltf* promoter displays altered expression of PCP, aquaporins, and EMT proteins at gestational day 3.5. (A) H&E stain of 6-month-old *Prickle1*$^{f/f}$ *Ltf*$^{+/icre}$ cKO compared to control at GD3.5. (B and C) Immunohistochemistry stain of 6-month-old *Prickle1*$^{f/f}$ *Ltf*$^{+/icre}$ cKO compared to control at GD3.5 for Krt7 (B) and Sox9 (C). (D-G) Immunofluorescence stain of 6-month-old *Prickle1*$^{f/f}$ *Ltf*$^{+/icre}$ cKO compared to control at GD3.5 for AQP2 (D), Zeb1 (E), Twist1 (F), and Snai2 (G). (H) Immunohistochemistry stain of 6-month-old *Prickle1*$^{f/f}$ *Ltf*$^{+/icre}$ cKO compared to control at GD3.5 for Vimentin. (Scale bars, 100 $\mu$m).

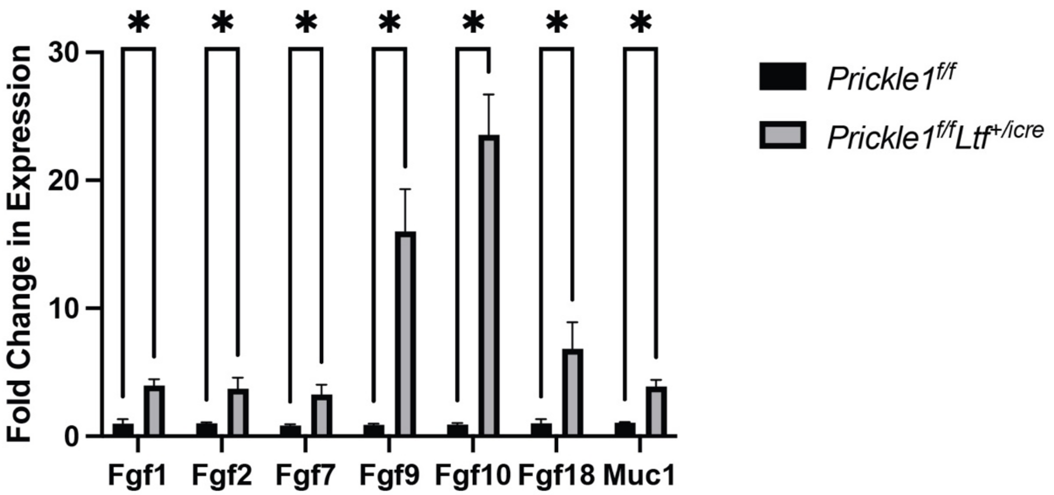


**Supplemental Figure 2.** Gene expression of fibroblast growth factors and *Muc-1* in 6-month-old *Prickle1$^{f/f}$ Ltf$^{+/icre}$* mice compared with control (n=3). Error bars represent ± SEM. Student's *t* test was performed, *$P$<0.05.

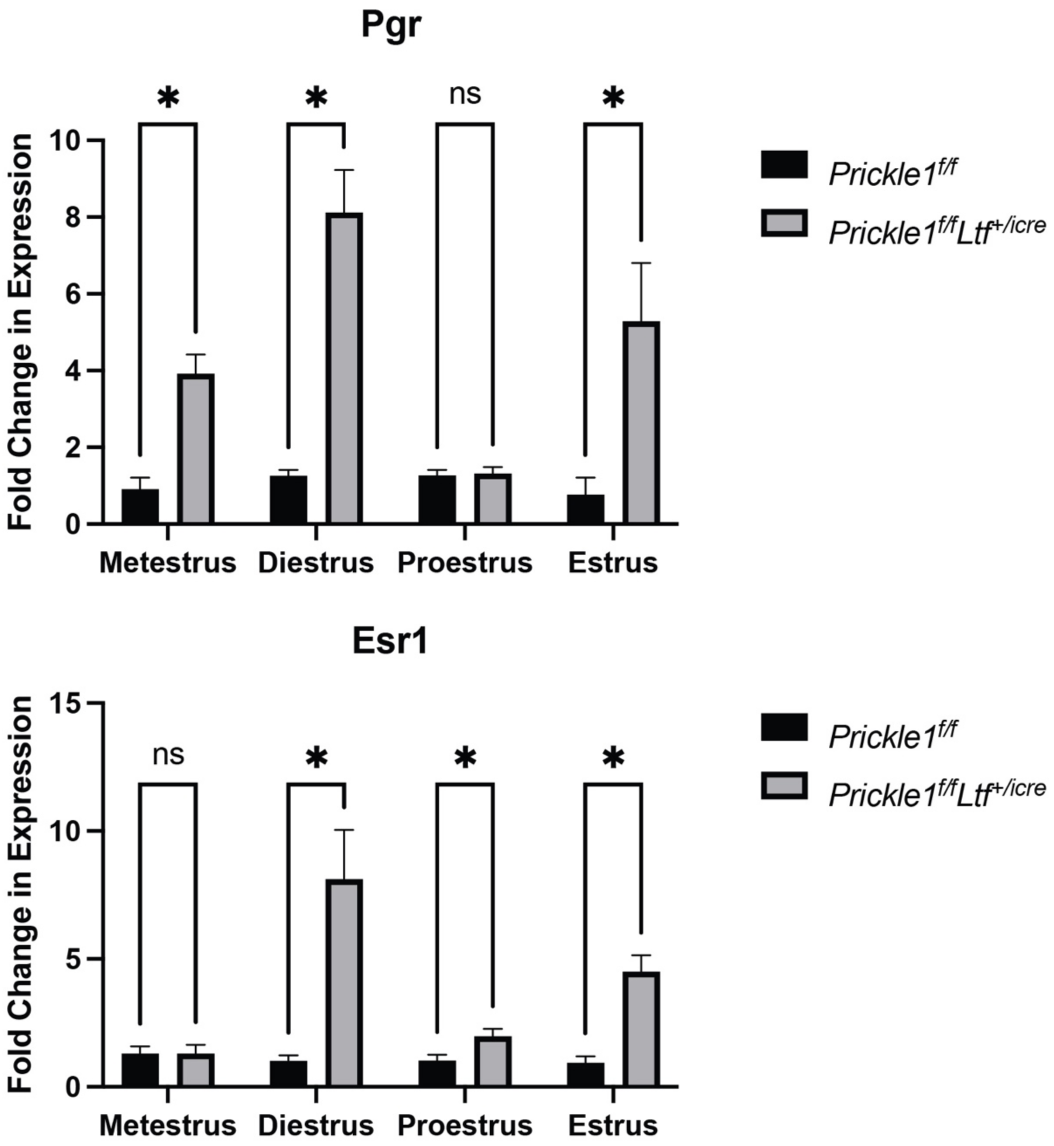


**Supplemental Figure 3.** Gene expression of *Pgr* and *Esr1* in 6-month-old *Prickle1*$^{f/f}$ *Ltf*$^{+/icre}$ mice compared with control (n=3) across estrus stages. Error bars represent ± SEM. Student's *t* test was performed, *$P$<0.05.

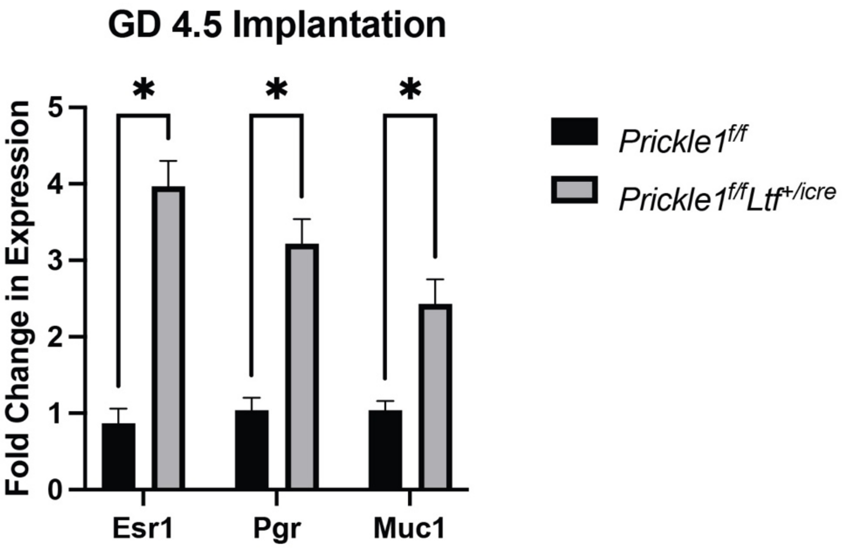


**Supplemental Figure 4.** Gene expression of *Esr1, Pgr,* and *Muc-1* in 6-month-old *Prickle1$^{f/f}$ Ltf$^{+/icre}$* mice compared with control (n=3) at gestational day 4.5. Error bars represent ± SEM. Student's *t* test was performed, **P*<0.05.

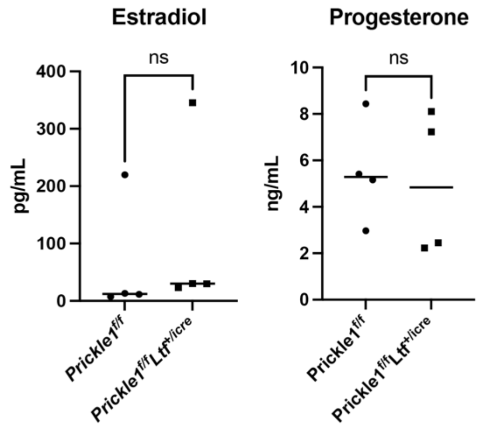


**Supplemental Figure 5.** Serum levels of estrogen and progesterone in 4-month-old *Prickle1$^{f/f}$ Ltf$^{+/icre}$* mice compared with control (n=4).

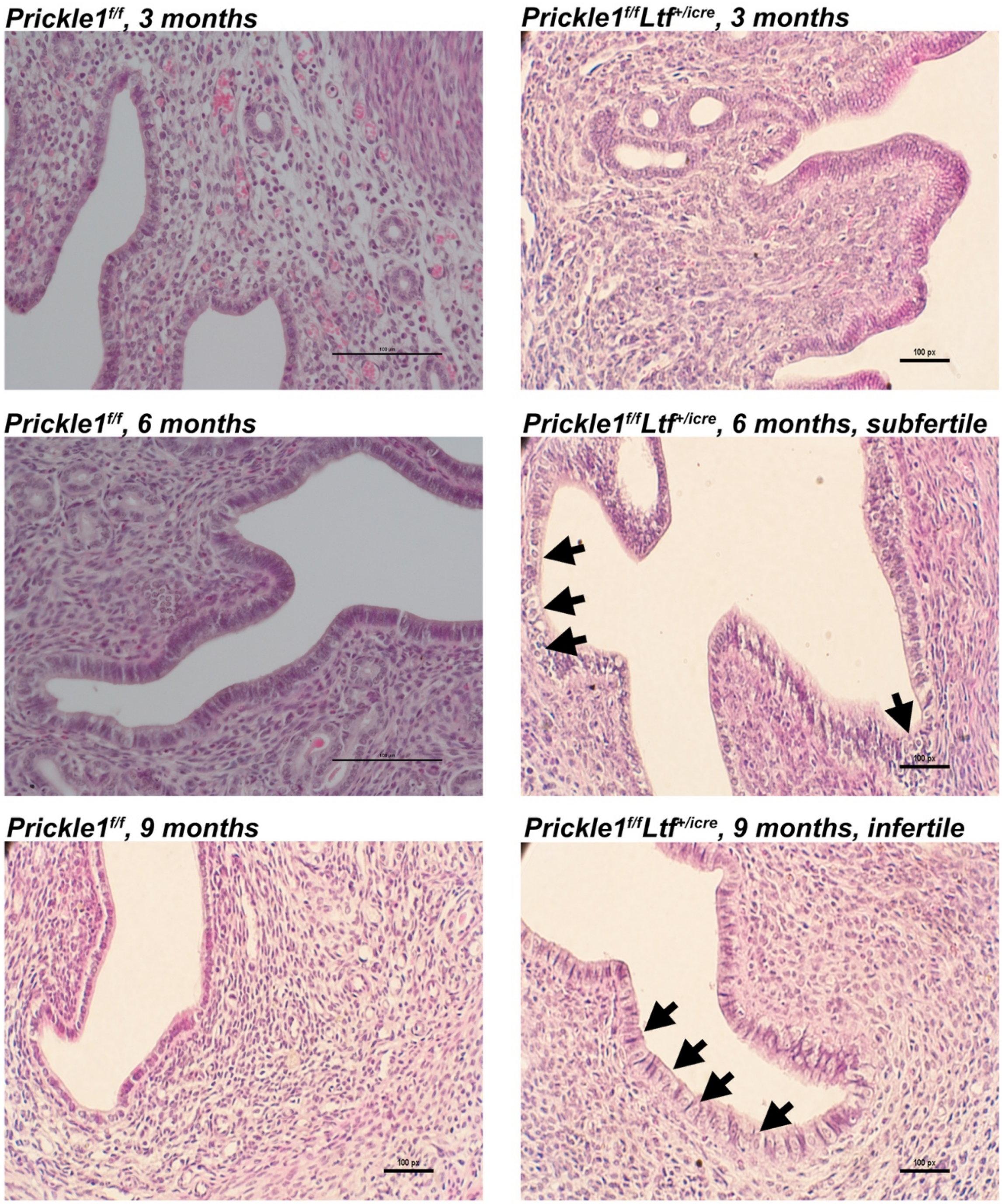


**Supplemental Figure 6.** <u>Characterization of uteri in *Prickle1$^{f/f}$ Ltf$^{+/icre}$* mice</u>. H&E stain of 3, 6, and 9-month-old *Prickle1$^{f/f}$ Ltf$^{+/icre}$* cKO compared to corresponding controls in similar estrus cycle stages with corresponding fertility information. Arrows indicate multinucleated cells. (Scale bars, 100 μm).

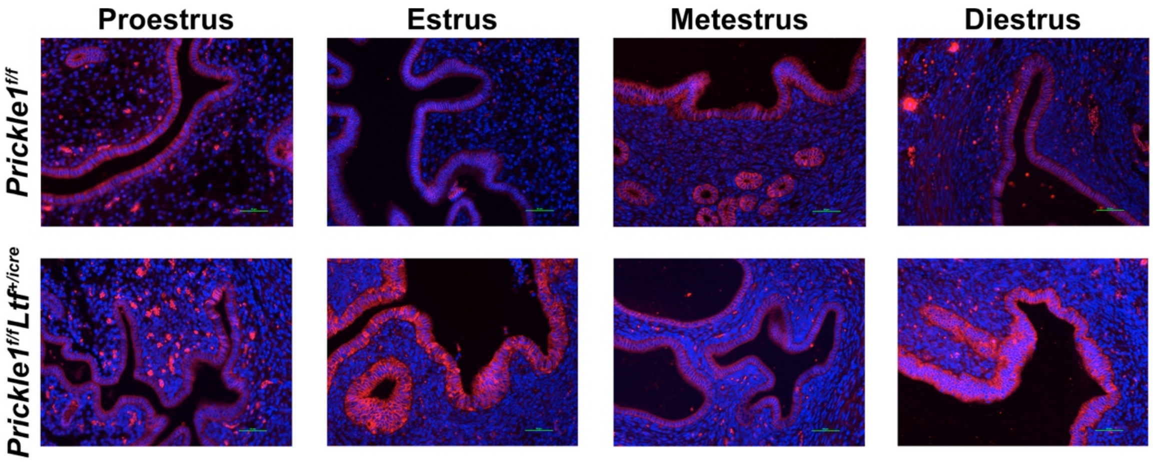


**Supplemental Figure 7.** E-cadherin expression throughout the estrus cycles. Immunofluorescence stain of 6-month-old *Prickle1*$^{f/f}$ *Ltf*$^{+/icre}$ cKO compared to control in various estrus cycle stages for epithelial marker E-cadherin (red) and DAPI (blue) indicating high baso-lateral and overall E-cadherin expression in mutant throughout the estrus cycle. (Scale bars, 50 $\mu$m).

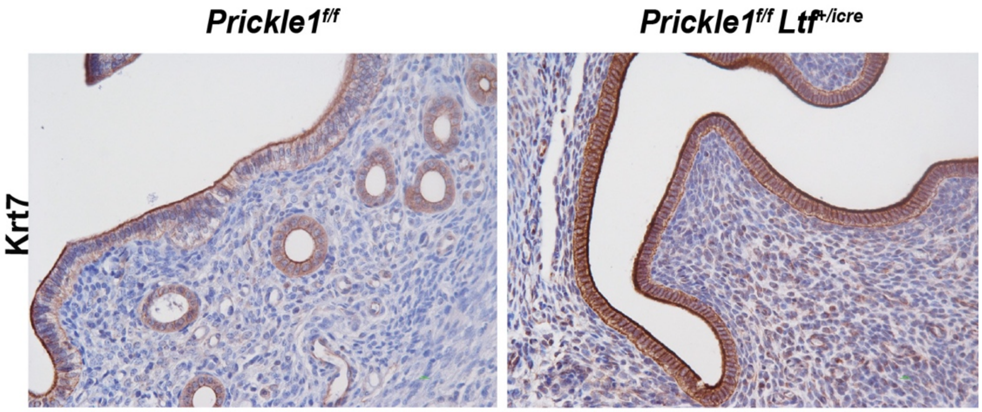


**Supplemental Figure 8.** Loss of *Prickle1* under the *Ltf* promoter alters cellular structure and dysregulates expression of Wnt/PCP proteins. Immunohistochemistry stain of 6-month-old *Prickle1*$^{f/f}$ *Ltf*$^{+/icre}$ cKO compared to control in diestrus for KRT7, indicating higher expression of KRT7 in mutant. (Scale bars, 10 $\mu$m).

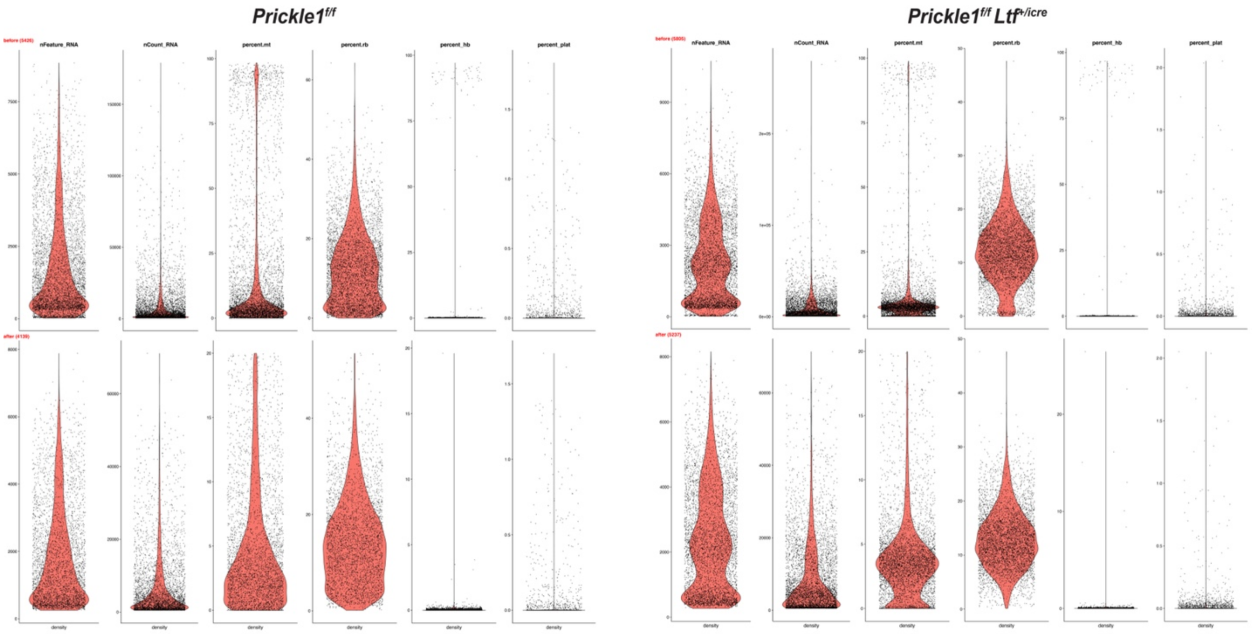


**Supplemental Figure 9.** Quality control distributions for single cell RNA sequencing. Violin plots depicting the number of features (nFeature_RNA), number of unique molecular identifiers (mCount_RNA), and percentage reads mapped to mitochondrial genes (percent.mt) in each cell, before and after QC filtering in control and *Prickle1*$^{f/f}$ *Ltf*$^{+/icre}$ uterine samples.

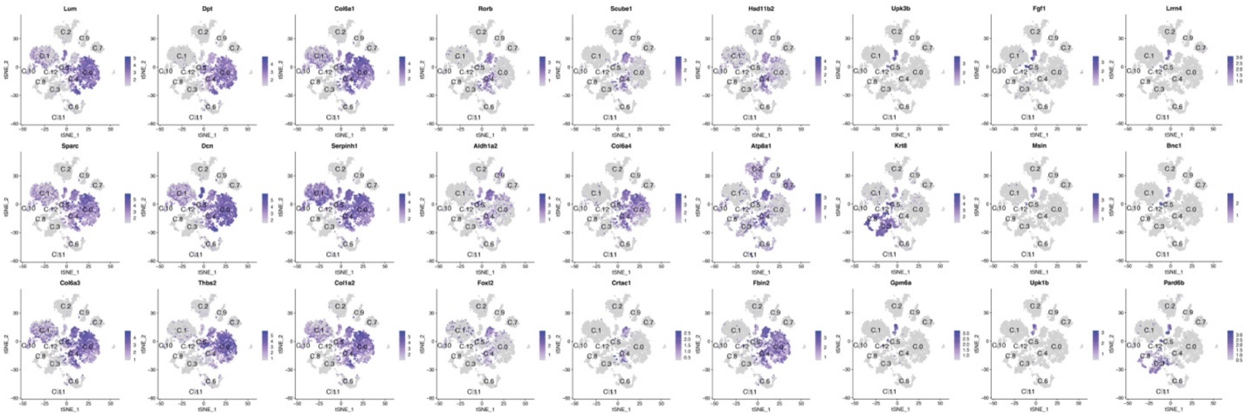


**Supplemental Figure 10.** Clusters of cells showing conserved markers of uterine stromal cells.

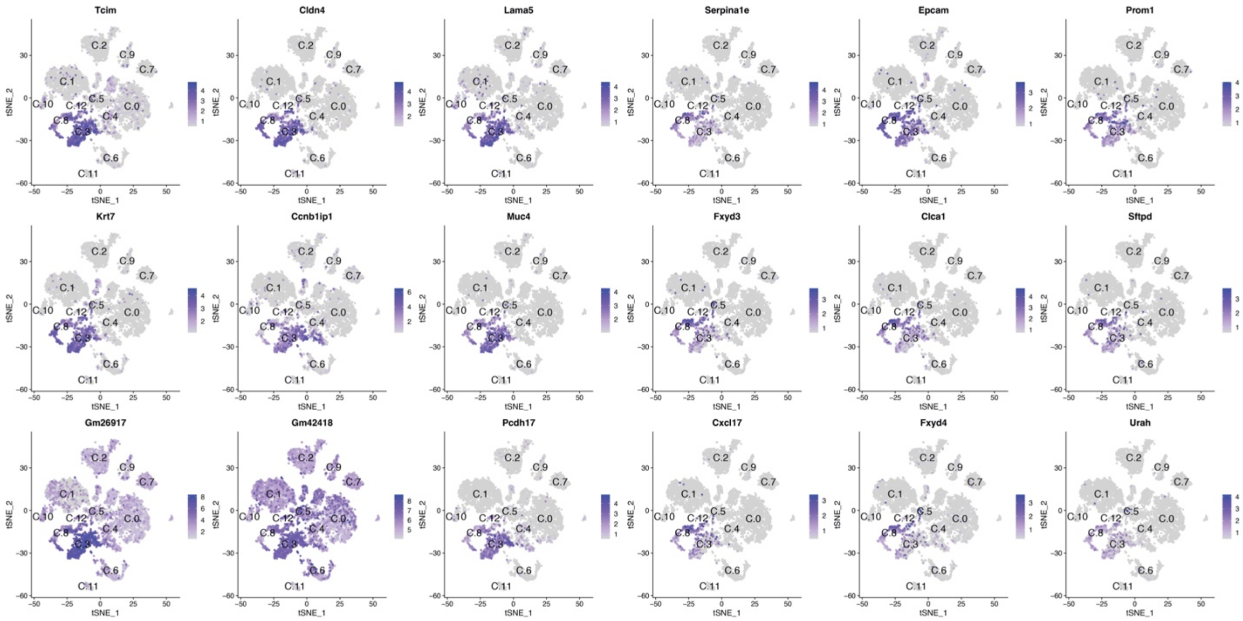


**Supplemental Figure 11.** <u>Clusters of cells showing conserved markers of uterine epithelial cells.</u>

A.

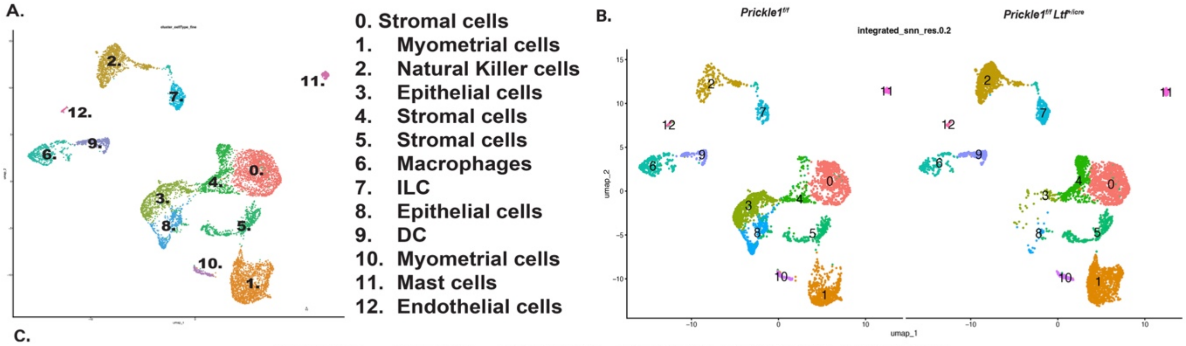


C.

| Cluster | Sample-1 | Sample-2 | Number of cells in Sample-1 | Number of cells in Sample-2 | Expected number of cells in Sample-1 | Expected number of cells in Sample-2 | Percentage deviation in Sample-1 | Percentage deviation in Sample-2 | Chi-square pvalue | Bonferroni pvalue |
|---|---|---|---|---|---|---|---|---|---|---|
| 0 | PK1_fl-fl | PK1_fl-fl_ltf-cre | 760 | 1090 | 816.676 | 1033.324 | -6.940 | 5.485 | 0.003057822 | **0.039751685** |
| 1 | PK1_fl-fl | PK1_fl-fl_ltf-cre | 598 | 1095 | 747.368 | 945.632 | -19.986 | 15.796 | 6.66134E-16 | **8.65974E-15** |
| 2 | PK1_fl-fl | PK1_fl-fl_ltf-cre | 217 | 879 | 483.825 | 612.175 | -55.149 | 43.586 | 0 | **0** |
| 3 | PK1_fl-fl | PK1_fl-fl_ltf-cre | 822 | 74 | 395.536 | 500.464 | 107.819 | -85.214 | 0 | **0** |
| 4 | PK1_fl-fl | PK1_fl-fl_ltf-cre | 198 | 657 | 377.437 | 477.563 | -47.541 | 37.573 | 0 | **0** |
| 5 | PK1_fl-fl | PK1_fl-fl_ltf-cre | 315 | 431 | 329.319 | 416.681 | -4.348 | 3.436 | 0.271133999 | 1 |
| 6 | PK1_fl-fl | PK1_fl-fl_ltf-cre | 432 | 247 | 299.742 | 379.258 | 44.124 | -34.873 | 0 | **0** |
| 7 | PK1_fl-fl | PK1_fl-fl_ltf-cre | 101 | 344 | 196.444 | 248.556 | -48.586 | 38.399 | 0 | **0** |
| 8 | PK1_fl-fl | PK1_fl-fl_ltf-cre | 397 | 33 | 189.822 | 240.178 | 109.143 | -86.260 | 0 | **0** |
| 9 | PK1_fl-fl | PK1_fl-fl_ltf-cre | 127 | 215 | 150.975 | 191.025 | -15.880 | 12.550 | 0.007820557 | 0.101667247 |
| 10 | PK1_fl-fl | PK1_fl-fl_ltf-cre | 72 | 78 | 66.217 | 83.783 | 8.734 | -6.902 | 0.337754116 | 1 |
| 11 | PK1_fl-fl | PK1_fl-fl_ltf-cre | 79 | 50 | 56.947 | 72.053 | 38.727 | -30.607 | 8.23418E-05 | **0.001070444** |
| 12 | PK1_fl-fl | PK1_fl-fl_ltf-cre | 21 | 44 | 28.694 | 36.306 | -26.814 | 21.192 | 0.053784449 | 0.69919784 |

**Supplemental Figure 12.** Loss of PRICKLE1 leads to changes in uterine cell clusters. (A) SingleR program predictions for cell clusters present in control and *Prickle1*$^{f/f}$ *Ltf*$^{+/icre}$ mice uteri shown by UMAP plot. Cell clusters were given a number between 0-12. (B) Individual UMAP plots for control and *Prickle1*$^{f/f}$ *Ltf*$^{+/icre}$ mice showing changes in cell clusters. (C) Table showing expected number of cells in each cluster compared to actual number of cells in each cluster. Percentage deviation, Chi-square p-value, and Bonferroni p-value show significant changes in clusters caused by loss of PRICKLE1 in *Prickle1*$^{f/f}$ *Ltf*$^{+/icre}$ mice.

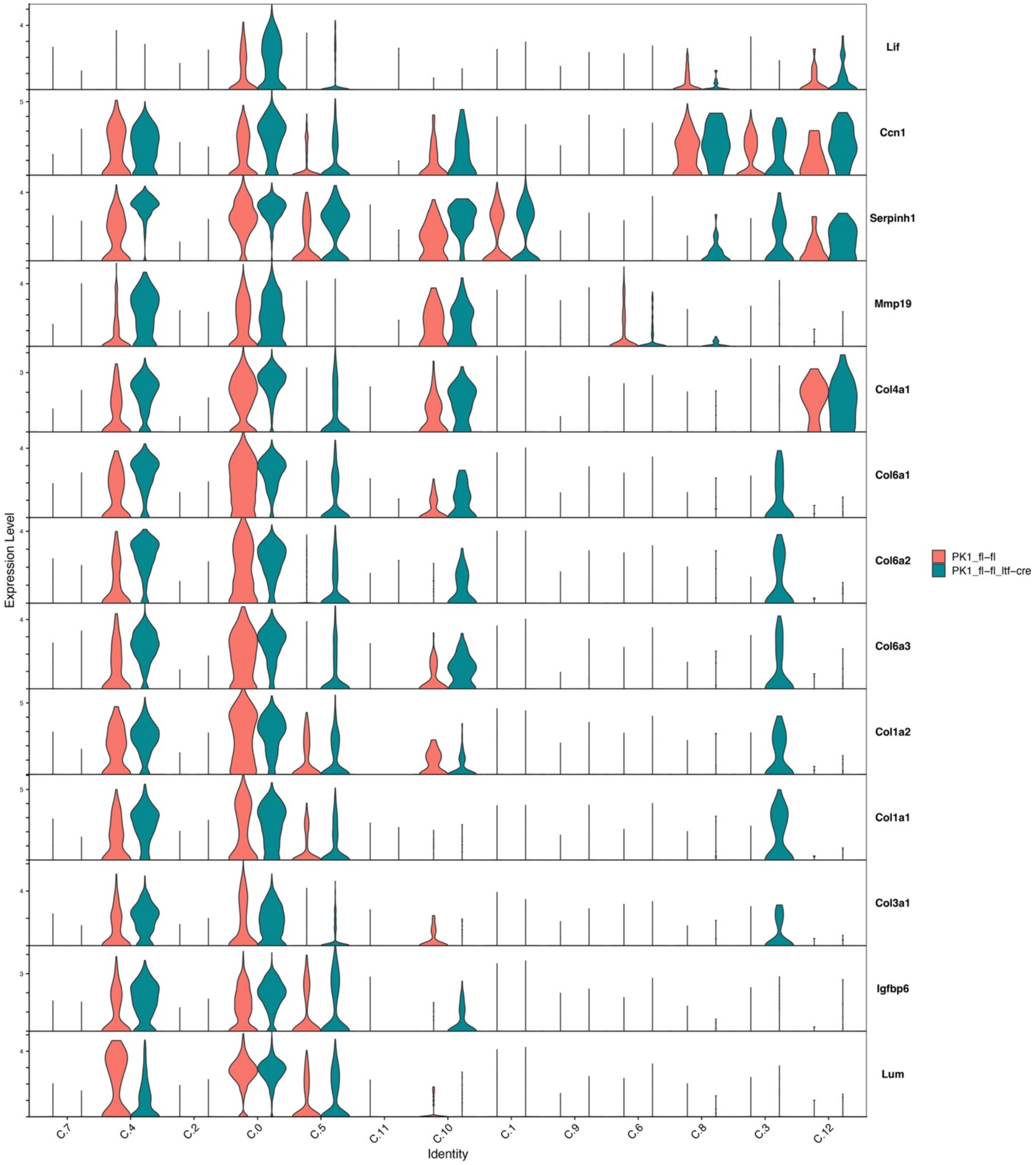


**Supplemental Figure 13.** Cluster expression of select collagens and REST target genes in *Prickle1$^{f/f}$ Ltf$^{+/icre}$* mice. Violin plots from single cell RNA sequencing showing expression (log 2-fold expression > 0) of collagens and REST target genes in different cell type clusters. X-axis represents the cluster number and Y-axis represents the gene expression level.

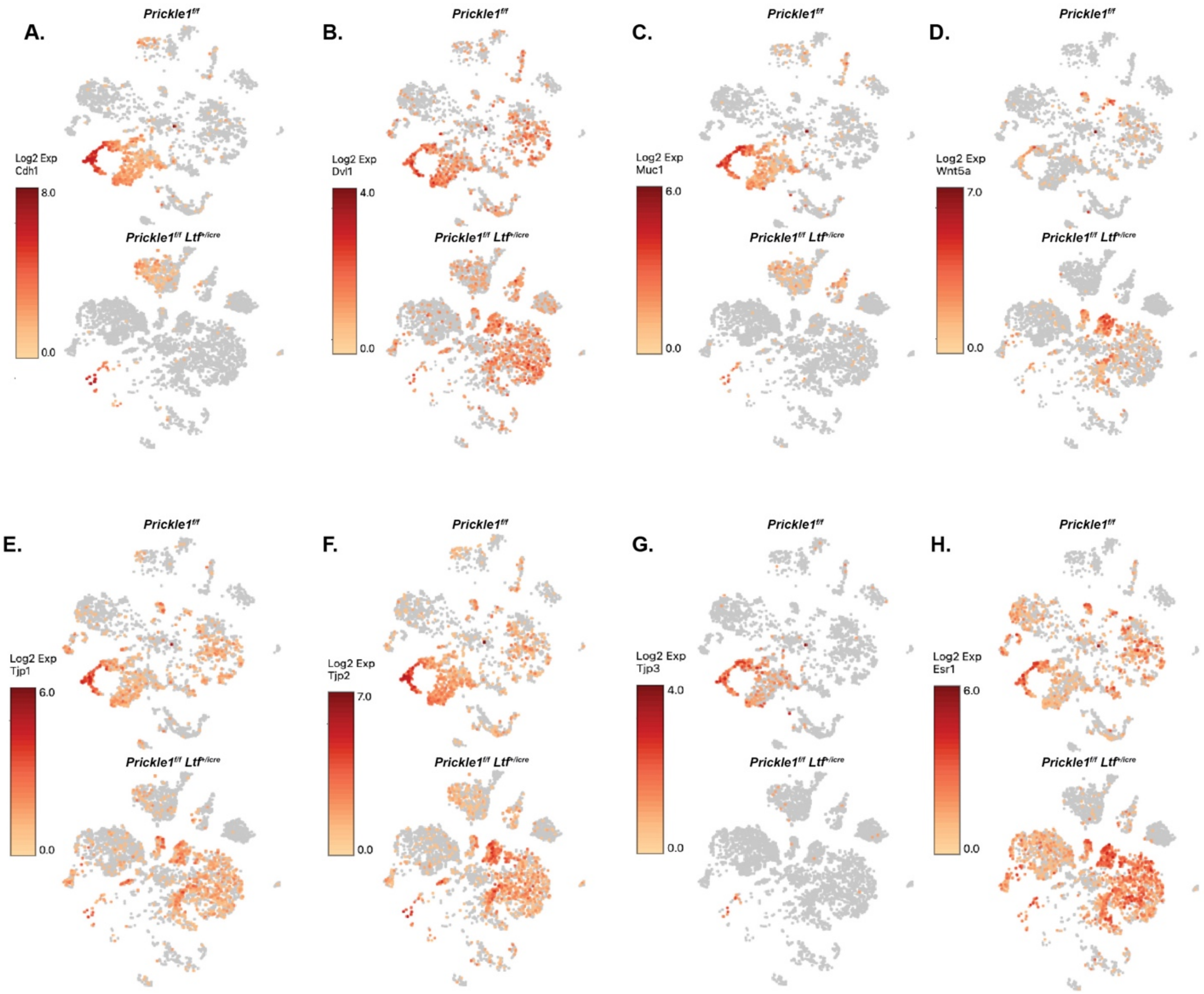


**Supplemental Figure 14.** Comparative t-SNE plots showing expression (log 2-fold expression > 0) of Wnt/PCP components (A) *Cdh1,* (B) *Dvl1*, (C) *Muc1*, (D) *Wnt5a*, (E) *Tjp1*, (F) *Tjp2*, (G) *Tjp3*, and (H) *Esr1*. Dark red indicates higher levels of expression.

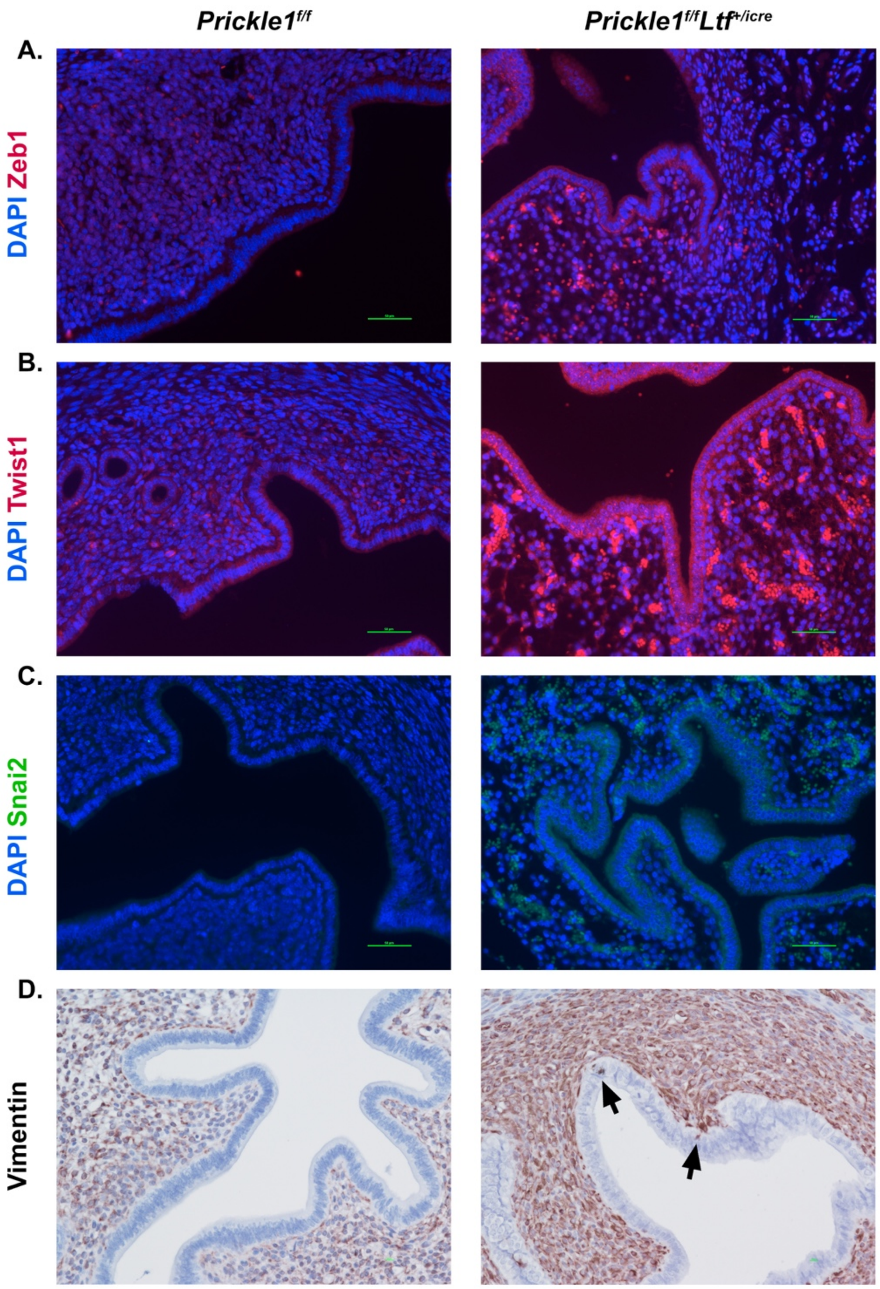


**Supplemental Figure 15.** Loss of *Prickle1* under the *Ltf* promoter displays increased expression of EMT proteins. (A) Immunofluorescence stain of 6-month-old *Prickle1*$^{f/f}$ *Ltf*$^{+/icre}$ cKO compared to control in diestrus for EMT marker ZEB1 (red) and DAPI (blue) indicating high ZEB1 expression in mutant epithelium. (Scale bars, 50 μm). (B) Immunofluorescence stain of 6-month-old *Prickle1*$^{f/f}$ *Ltf*$^{+/icre}$ cKO compared to control in diestrus for EMT marker TWIST1 (red) and DAPI (blue) indicating high TWIST1 expression in mutant epithelium. (Scale bars, 50 μm). (C) Immunofluorescence stain of 6-month-old *Prickle1*$^{f/f}$ *Ltf*$^{+/icre}$ cKO compared to control in diestrus for EMT marker SNAI2 (green) and DAPI (blue) indicating high SNAI2 expression in mutant epithelium. (Scale bars, 50 μm). (D) Immunohistochemistry staining of 6-month-old *Prickle1*$^{f/f}$ *Ltf*$^{+/icre}$ cKO compared to control in diestrus for Vimentin indicating higher vimentin expression in the stroma. Arrows indicate epithelial cells that are stained positive for vimentin. (Scale bars, 10 μm).

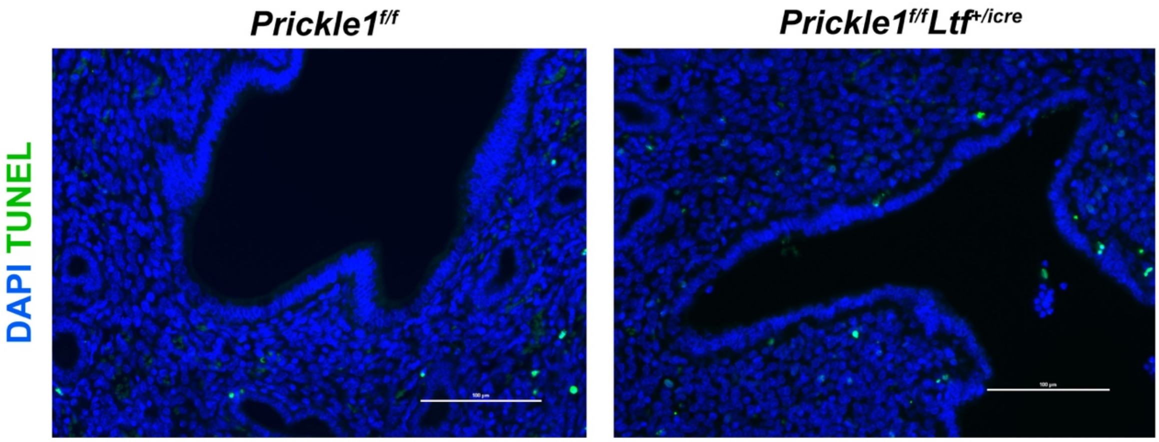


**Supplemental Figure 16.** Loss of *Prickle1* under the *Ltf* promoter does not increase apoptosis. (A) Immunofluorescence stain of 6-month-old control and *Prickle1*$^{f/f}$ *Ltf*$^{+/icre}$ cKO in diestrus for apoptosis marker TUNEL (green) and DAPI (blue). (Scale bars, 100 μm).

**Supplemental Table 1.** Pathway analysis of predicted physiological system development and functions associated with dysregulated genes in the epithelial cell population (cluster 3) of *Prickle1*$^{f/f}$ *Ltf*$^{+/icre}$ cKO mice.

| **Physiological System Development and Function** | ***P* value** |
|---|---|
| Organismal survival | 7.28E-46 |
| Cardiovascular system development and function | 1.70E-42 |
| Organismal development | 1.70E-42 |
| Embryonic development | 7.43E-40 |
| Tissue development | 3.94E-33 |

Function predictions made by IPA software based off genes which were found to be dysregulated in epithelial cells of 6-month-old *Prickle1*$^{f/f}$ *Ltf*$^{+/icre}$ mice cKO (n=3) in diestrus.

**Supplemental Table 2.** Pathway analysis of predicted disease and disorders associated with dysregulated genes in the epithelial cell population (cluster 8) of *Prickle1*$^{f/f}$ *Ltf*$^{+/icre}$ cKO mice.

| **Disease and Disorders** | ***P* value** |
|---|---|
| Cancer | 2.30E-45 |
| Organismal injury and abnormalities | 2.30E-45 |
| Endocrine system disorder | 3.14E-39 |
| Gastrointestinal disease | 4.80E-36 |
| Respiratory disease | 1.69E-17 |

Disease predictions made by IPA software based off genes which were found to be dysregulated in epithelial cells of 6-month-old *Prickle1*$^{f/f}$ *Ltf*$^{+/icre}$ mice cKO (n=3) in diestrus.

**Supplemental Table 3.** Pathway analysis of predicted molecular and cellular functions associated with dysregulated genes in the epithelial cell population (cluster 8) of *Prickle1*$^{f/f}$ *Ltf*$^{+/icre}$ cKO mice.

| **Molecular and Cellular Functions** | ***P* value** |
|---|---|
| Cell death and survival | 1.22E-18 |
| Cellular function and maintenance | 2.10E-16 |
| Cellular development | 2.42E-12 |
| Cellular growth and proliferation | 2.42E-12 |
| DNA replication, recombination, and repair | 1.44E-11 |

Function predictions made by IPA software based off genes which were found to be dysregulated in epithelial cells of 6-month-old *Prickle1*$^{f/f}$ *Ltf*$^{+/icre}$ mice cKO (n=3) in diestrus.

**Supplemental Table 4.** Pathway analysis of predicted physiological system development and functions associated with dysregulated genes in the epithelial cell population (cluster 8) of *Prickle1*$^{f/f}$ *Ltf*$^{+/icre}$ cKO mice.

| **Physiological System Development and Function** | ***P* value** |
|---|---|
| Organismal survival | 5.73E-17 |
| Hair and skin development and function | 5.32E-10 |
| Embryonic development | 1.68E-08 |
| Organismal development | 1.73E-08 |
| Connective tissue development and function | 3.16E-08 |

Function predictions made by IPA software based off genes which were found to be dysregulated in epithelial cells of 6-month-old *Prickle1*$^{f/f}$ *Ltf*$^{+/icre}$ mice cKO (n=3) in diestrus.

**Supplemental Table 5.** Pathway analysis of predicted disease and disorders associated with dysregulated genes in the stromal cell population (cluster 0) of *Prickle1*$^{f/f}$ *Ltf*$^{+/icre}$ cKO mice.

| Disease and Disorders | *P* value |
|---|---|
| Connective tissue disorders | 1.08E-21 |
| Inflammatory disease | 1.08E-21 |
| Organismal injury and abnormalities | 1.08E-21 |
| Skeletal and muscular disorders | 1.08E-21 |
| Inflammatory response | 1.00E-19 |

Disease predictions made by IPA software based off genes which were found to be dysregulated in stromal cells of 6-month-old *Prickle1*$^{f/f}$ *Ltf*$^{+/icre}$ mice cKO (n=3) in diestrus.

**Supplemental Table 6.** Pathway analysis of predicted molecular and cellular functions associated with dysregulated genes in the stromal cell population (cluster 0) of *Prickle1*$^{f/f}$ *Ltf*$^{+/icre}$ cKO mice.

| **Molecular and Cellular Functions** | ***P* value** |
|---|---|
| Cellular movement | 4.63E-18 |
| Cell-to-cell signaling and interaction | 1.73E-11 |
| Cellular development | 1.01E-10 |
| Cellular growth and proliferation | 1.01E-10 |
| Cell death and survival | 2.37E-09 |

Function predictions made by IPA software based off genes which were found to be dysregulated in stromal cells of 6-month-old *Prickle1*$^{f/f}$ *Ltf*$^{+/icre}$ mice cKO (n=3) in diestrus.

**Supplemental Table 7.** Pathway analysis of predicted physiological system development and functions associated with dysregulated genes in the stromal cell population (cluster 0) of *Prickle1*$^{f/f}$ *Ltf*$^{+/icre}$ cKO mice.

| **Physiological System Development and Function** | ***P* value** |
|---|---|
| Hematological system development and function | 4.63E-18 |
| Immune cell trafficking | 4.63E-18 |
| Tissue morphology | 7.04E-13 |
| Cardiovascular system development and function | 8.64E-11 |
| Organismal development | 8.64E-11 |

Function predictions made by IPA software based off genes which were found to be dysregulated in stromal cells of 6-month-old *Prickle1*$^{f/f}$ *Ltf*$^{+/icre}$ mice cKO (n=3) in diestrus.

**Supplemental Table 8.** Pathway analysis of predicted molecular and cellular functions associated with dysregulated genes in the stromal cell population (cluster 4) of *Prickle1*$^{f/f}$ *Ltf*$^{+/icre}$ cKO mice.

| Molecular and Cellular Functions | *P* value |
|---|---|
| Cell death and survival | 3.84E-13 |
| Cellular development | 1.17E-10 |
| Cellular growth and proliferation | 1.17E-10 |
| Cellular movement | 3.59E-10 |
| Amino acid metabolism | 3.16E-09 |

Function predictions made by IPA software based off genes which were found to be dysregulated in stromal cells of 6-month-old *Prickle1*$^{f/f}$ *Ltf*$^{+/icre}$ mice cKO (n=3) in diestrus.

**Supplemental Table 9.** Pathway analysis of predicted physiological system development and functions associated with dysregulated genes in the stromal cell population (cluster 4) of *Prickle1*$^{f/f}$ *Ltf*$^{+/icre}$ cKO mice.

| **Physiological System Development and Function** | ***P* value** |
|---|---|
| Organismal survival | 8.67E-12 |
| Embryonic development | 2.30E-11 |
| Organismal development | 2.30E-11 |
| Reproductive system development and function | 1.63E-08 |
| Connective tissue development and function | 2.58E-08 |

Function predictions made by IPA software based off genes which were found to be dysregulated in stromal cells of 6-month-old *Prickle1*$^{f/f}$ *Ltf*$^{+/icre}$ mice cKO (n=3) in diestrus.

**Supplemental Table 10.** Pathway analysis of predicted disease and disorders associated with dysregulated genes in the stromal cell population (cluster 5) of *Prickle1*$^{f/f}$ *Ltf*$^{+/icre}$ cKO mice.

| **Disease and Disorders** | ***P* value** |
|---|---|
| Inflammatory response | 2.51E-20 |
| Organismal injury and abnormalities | 2.51E-20 |
| Cancer | 3.18E-14 |
| Dermatological diseases and conditions | 7.38E-14 |
| Inflammatory disease | 7.38E-14 |

Disease predictions made by IPA software based off genes which were found to be dysregulated in stromal cells of 6-month-old *Prickle1*$^{f/f}$ *Ltf*$^{+/icre}$ mice cKO (n=3) in diestrus.

**Supplemental Table 11.** Pathway analysis of predicted molecular and cellular functions associated with dysregulated genes in the stromal cell population (cluster 5) of *Prickle1*$^{f/f}$ *Ltf*$^{+/icre}$ cKO mice.

| **Molecular and Cellular Functions** | ***P* value** |
|---|---|
| Cellular movement | 2.64E-20 |
| Cell-to-cell signaling and interaction | 7.17E-18 |
| Cellular growth and proliferation | 7.88E-13 |
| Cell death and survival | 1.08E-12 |
| Cellular development | 3.26E-12 |

Function predictions made by IPA software based off genes which were found to be dysregulated in stromal cells of 6-month-old *Prickle1*$^{f/f}$ *Ltf*$^{+/icre}$ mice cKO (n=3) in diestrus.

**Supplemental Table 12.** Pathway analysis of predicted physiological system development and functions associated with dysregulated genes in the stromal cell population (cluster 5) of *Prickle1*$^{f/f}$ *Ltf*$^{+/icre}$ cKO mice.

| **Physiological System Development and Function** | ***P* value** |
|---|---|
| Cardiovascular system development and function | 1.07E-14 |
| Immune cell trafficking | 3.36E-14 |
| Hematological system development and function | 5.37E-14 |
| Organismal development | 4.56E-13 |
| Organismal survival | 6.01E-13 |

Function predictions made by IPA software based off genes which were found to be dysregulated in stromal cells of 6-month-old *Prickle1*$^{f/f}$ *Ltf*$^{+/icre}$ mice cKO (n=3) in diestrus.

**Supplemental Table 13.** Pathway analysis of predicted diseases and disorders associated with dysregulated genes in the natural killer cell population (cluster 2) of *Prickle1*$^{f/f}$ *Ltf*$^{+/icre}$ cKO mice.

| Disease or Disorder | *P* value |
|---|---|
| Immunological disease | 6.14E-30 |
| Organismal injury and abnormalities | 6.14E-30 |
| Cancer | 4.20E-25 |
| Hematological disease | 4.20E-25 |
| Inflammatory disease | 6.36E-24 |

Disease predictions made by IPA software based off genes which were found to be dysregulated in natural killer cells of 6-month-old *Prickle1*$^{f/f}$ *Ltf*$^{+/icre}$ mice cKO (n=3) in diestrus.

**Supplemental Table 14.** Pathway analysis of predicted diseases and disorders associated with dysregulated genes in the natural killer cell population (cluster 2) of *Prickle1*$^{f/f}$ *Ltf*$^{+/icre}$ cKO mice.

| **Molecular and Cellular Functions** | ***P* value** |
|---|---|
| Cell cycle | 1.15E-20 |
| Cellular assembly and organization | 4.60E-20 |
| DNA replication, recombination, and repair | 4.60E-20 |
| Cellular growth and proliferation | 3.43E-17 |
| Cellular development | 1.55E-14 |

Function predictions made by IPA software based off genes which were found to be dysregulated in natural killer cells of 6-month-old *Prickle1*$^{f/f}$ *Ltf*$^{+/icre}$ mice cKO (n=3) in diestrus.

**Supplemental Table 15.** Pathway analysis of predicted diseases and disorders associated with dysregulated genes in the natural killer cell population (cluster 2) of *Prickle1*$^{f/f}$ *Ltf*$^{+/icre}$ cKO mice.

| **Physiological System Development and Function** | ***P* value** |
|---|---|
| Lymphoid tissue structure and development | 3.43E-17 |
| Hematological system development and function | 1.55E-14 |
| Organismal survival | 5.15E-13 |
| Hematopoiesis | 1.55E-14 |
| Tissue development | 1.55E-14 |

Function predictions made by IPA software based off genes which were found to be dysregulated in natural killer cells of 6-month-old *Prickle1*$^{f/f}$ *Ltf*$^{+/icre}$ mice cKO (n=3) in diestrus.

**Supplemental Table 16.** The number of proper and improperly oriented dividing cells in *Prickle1*$^{f/f}$ *Ltf*$^{+/icre}$ and control mice. (n=9 sections each).

| | **Proper Orientation** | **Improperly Orientation** |
|---|---|---|
| *Prickle1*$^{f/f}$ | 19 | 3 |
| *Prickle1*$^{f/f}$ *Ltf*$^{+/icre}$ | 5 | 13 |

**Supplemental Table 17.** Numbers of singlets and doublets within the single-cell RNA sequencing data for *Prickle1*$^{f/f}$ *Ltf*$^{+/icre}$ and control mice.

| | **Singletons** | **Doublets** | **Row Totals** |
|---|---|---|---|
| *Prickle1*$^{f/f}$ | 5289 (52.64) [0.11] | 137 (161.36) [3.68] | 5426 |
| *Prickle1*$^{f/f}$ *Ltf*$^{+/icre}$ | 5608 (5632.36) [0.11] | 197 (172.64) [3.44] | 5805 |
| **Column Totals** | 10897 | 334 | **11231 (Grand Total)** |

** the observed cell totals, (the expected cell totals) and [the chi-square statistic for each cell]

The chi-square statistic is 7.3355. The p-value is .00676. The result is significant at $p < .05$.

.
